# SUMO paralogues differentially affect phase separation and aggregation of intrinsically disordered proteins

**DOI:** 10.64898/2026.05.11.724321

**Authors:** Philipp Schönberger, Supriyo Naskar, Oliver Ordowski, Joshua Vollrath, Jia-Xuan Chen, Martin M. Möckel, Márton Gelléri, Christian Renz, Julian D. Langer, Kurt Kremer, Oleksandra Kukharenko, Helle D. Ulrich

## Abstract

SUMO, the small ubiquitin-like modifier, modulates interactions of folded proteins, but also affects the dynamics of biomolecular condensates. Moreover, SUMO can directly impinge on the biophysical properties of its targets, often increasing their solubility. TDP-43 is an RNA-binding protein containing an intrinsically disordered domain that participates in protein phase separation. Its aggregation, linked to neurodegenerative diseases, is counteracted by SUMOylation, but the underlying principles are poorly understood. We now reveal distinct mechanisms of how the two isoforms, SUMO1 and SUMO2, act on TDP-43. While SUMO1 inhibits phase separation by blocking TDP-43’s self-assembly site, SUMO2 enhances phase separation via proximity-induced self-interaction involving its N-terminus and promotes a liquid-like state that prevents aggregation. The latter represents a general principle by which SUMO solubilises other targets. Our findings provide insight into how SUMOylation regulates transitions between soluble, liquid-like and aggregated states.

## Introduction

Protein phase separation has emerged as an organising principle of many membraneless organelles, including the nucleolus, PML nuclear bodies, and cytoplasmic stress granules^1^. Their cohesion often depends on proteins containing intrinsically disordered regions (IDRs) that engage in dynamic, multivalent interactions^2^. These facilitate biological reactions by generating protein-enriched microenvironments. At the same time, the high local concentration of unstructured peptides within such biomolecular condensates can promote a transition to insoluble aggregates, a process that has been implicated in several neurodegenerative diseases^3^. For example, under physiological conditions, the Transactive Response DNA Binding Protein 43 kDa (TDP-43), a nuclear RNA-binding protein, engages in phase separation via its N-terminal oligomerisation domain and a C-terminal low-complexity domain (LCD)^4–6^. In response to oxidative and other proteotoxic stress, TDP-43 can form cytoplasmic aggregates that are hallmarks of diseases like amyotrophic lateral sclerosis (ALS) and frontotemporal dementia (FTD)^7^. TDP-43’s LCD has been described as the major driver of aggregation, exhibiting prion-like behaviour^8–10^, and liquid-liquid phase separation is considered an intermediate step in fibril formation^11^.

Protein phase separation and aggregation are frequently modulated by posttranslational modifications (PTMs), ranging from phosphorylation and acetylation to ubiquitylation and SUMOylation^12^. SUMO, the small ubiquitin-like modifier, exists in several isoforms. Among these, SUMO1 shares less than 50% identity throughout its sequence with the highly related SUMO2 and SUMO3, which are 95% identical and usually indistinguishable^13^. SUMO2/3 in particular is associated with cellular stress responses^14,15^. Regulation of protein-protein interactions by SUMO is mostly mediated by conserved SUMO interaction motifs (SIMs) that involve an extended peptide aligning with SUMO’s β-sheet in a parallel or antiparallel orientation^16,17^. Beyond the well-defined interactions of SUMOylated proteins with specific downstream effectors, SUMO is found to cover the surfaces of protein complexes or subcellular structures to produce larger assemblies in a glue-like fashion, a process coined group SUMOylation^18,19^. Multivalent SUMO-SIM contacts also form the basis of biomolecular condensates like PML nuclear bodies, which are held together by a network of dynamic interactions, facilitating the inclusion of other SUMO-binding proteins and at the same time functioning as a SUMOylation hub^20,21^. Similarly, many phase-separating proteins have been identified as SUMOylation targets^22,23^. Finally, SUMO apparently has the ability to affect the biophysical properties of its targets directly, in many cases enhancing their solubility, which has resulted in its frequent use as a purification tag^24^. Based on this notion, it is likely that SUMO impinges on protein phase separation and aggregation not only via SIM-dependent interactions, but also via intrinsic, SIM-independent effects.

According to proteomics-based screens, TDP-43 is SUMOylated at several sites^22,23,25^, and recent studies have confirmed modification by both SUMO1 and SUMO2/3^26–29^. However, the consequences of SUMOylation are still a matter of debate, and indirect effects have not always been excluded. Mutation of a conserved SUMOylation site in one of its RNA-binding domains was found to affect the subcellular localisation of TDP-43 and its aggregates and suggested that SUMO1 promotes TDP-43 aggregation^28^. In contrast, modification by SUMO2/3 was reported to protect TDP-43 from aggregation, both in association with PML nuclear bodies^26^ and in cytoplasmic stress granules^27^. The underlying mechanisms remain largely unexplored.

We have now systematically investigated the effects of SUMO1 and SUMO2 on the phase separation and aggregation of IDRs, using TDP-43’s LCD as a model case. Surprisingly, we found striking differences between the two isoforms in their effects on TDP-43^LCD^: while SUMO1 inhibited phase separation, SUMO2 rendered condensates more liquid and prevented the formation of solid aggregates. A combination of mass-spectrometry-based structural biology approaches and molecular dynamics (MD) simulations revealed distinct mechanisms underlying these two opposing phenotypes, relying primarily on intra- *versus* intermolecular interactions involving SUMO’s unstructured N-termini. By expanding our studies to a set of other IDRs with different characteristics, we provide evidence that the effect of SUMO2 is a general principle by which the modifier enhances the solubility of its target proteins. These findings not only demonstrate a complex role of isoform-specific SUMOylation in the physiology of TDP-43 and potentially other phase-separating proteins, but also suggest a general, intrinsic mechanism by which SUMO impinges on the biophysical properties of its targets.

## Results

### SUMO1 and SUMO2 have opposite effects on TDP-43^LCD^ in vitro

At room temperature and physiological salt concentration, the LCD of TDP-43 (**Fig. 1a**, amino acids 266-414) spontaneously undergoes phase separation *in vitro*^11^. Solubilisation and purification under native conditions is achieved by fusion to maltose-binding protein (MBP)^30^. After proteolytic cleavage of the C-terminal MBP via TEV protease, a fluorescently labelled TDP-43^LCD^ construct at a concentration of 50 µM separated into small droplets, as previously observed^11,30^, which slowly transitioned into a diffuse meshwork (**Fig. 1b**). A time course of turbidity measurements confirmed phase separation within 30 min (**Fig. 1c**). The presence of free SUMO1 or SUMO2 at equimolar concentration had no impact on droplet formation or turbidity (**Fig. S1a-c**).

**Fig 1:**
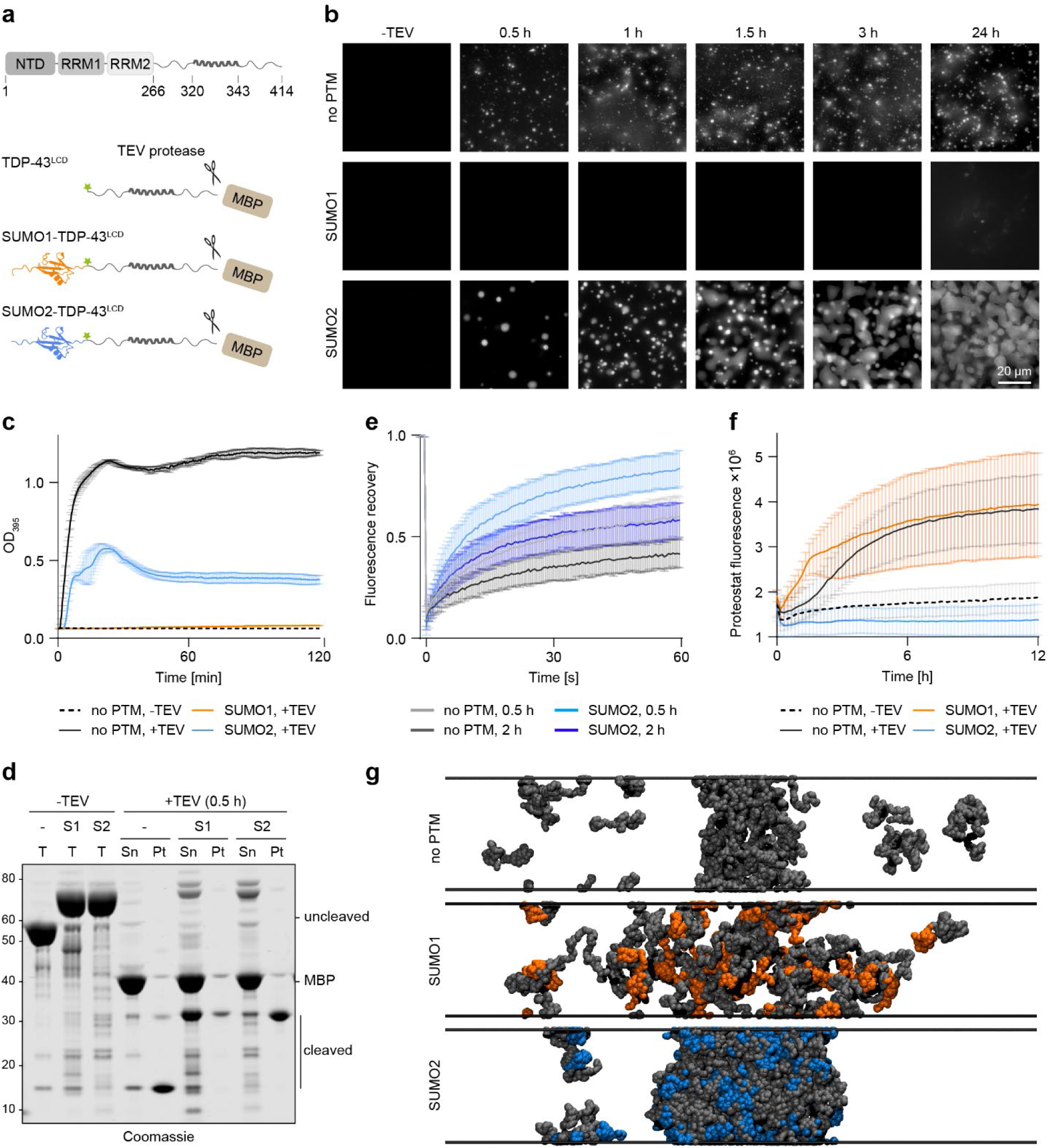
SUMO1 inhibits TDP-43^LCD^ phase separation *in vitro*, whereas SUMO2 renders condensates more liquid and prevents aggregation. **a,** Schematics of TDP-43 domain arrangement and constructs used for *in vitro* assays (NTD: N-terminal domain; RRM: RNA recognition motif; LCD: low-complexity domain; MBP: Maltose-binding protein, cleavable by TEV protease; green star: fluorescent label AzDye488). **b,** Covalent fusion of SUMO1 inhibits phase separation of TDP-43^LCD^, while SUMO2 enlarges droplet size. Condensate formation of TDP-43^LCD^ (50 µM, 5% AzDye488-labelled) fused to SUMO1 or SUMO2 as indicated, monitored by fluorescence microscopy in a time course after addition of TEV protease. **c,** Turbidity measurements of TDP-43^LCD^ constructs (50 µM) after addition of TEV protease. Values represent means ±SD of three technical replicates (one of three independent experiments). TEV cleavage patterns are shown in Fig. S1c. **d,** Sedimentation of TDP-43^LCD^ constructs (50 µM), analysed by SDS-PAGE and Coomassie staining following centrifugation 1 h after addition of TEV protease (T: total protein before addition of TEV protease; Sn: supernatant; P: pellet) (one of three independent experiments). Quantification is shown in Fig S1e. **e,** SUMO2 renders condensates more liquid. FRAP assays of TDP-43^LCD^ constructs (50 µM, 5% AzDye488-labelled), performed at indicated times after addition of TEV protease. Values represent means ±SD of approximately 10 bleached droplets (one of two independent experiments). **f,** SUMO2 prevents aggregation of TDP-43^LCD^. Aggregation of TDP-43^LCD^ constructs (50 µM, induced by 1% pre-aggregated protein), measured by Proteostat fluorescence. Lines represent means ±SD of 12 technical replicates performed in four measurements. **g,** Fusion of SUMO1 *versus* SUMO2 exerts opposite effects on TDP-43^LCD^ in MD simulations. Representative snapshots from CG slab coexistence MD simulations with the CALVADOS2 model at 300 K for unmodified TDP-43^LCD^, SUMO1–TDP-43^LCD^, and SUMO2–TDP-43^LCD^. Each system was composed of 100 chains and simulated for 5 μs (SUMO1: orange; SUMO2: blue; TDP-43^LCD^: dark grey).

To mimic SUMOylation, we followed an established strategy^31^ of fusing the modifier to the N-terminus of TDP-43^LCD^ (**Fig. 1a**). A mutation of SUMO’s C-terminal glycine (G) to valine (V) was introduced to enhance stability. Surprisingly, in this covalent arrangement, SUMO1 completely abolished droplet formation (**Fig. 1b**), even at protein concentrations up to 400 µM (**Fig. S1d**). Turbidity measurements showed no evidence of phase separation either (**Fig. 1c, S1c**). In stark contrast, SUMO2-TDP43^LCD^ readily formed extremely large condensates covering the bottom of the microscope slide (**Fig. 1b**), and turbidity measurements indicated rapid phase separation (**Fig. 1c, S1c**). In these assays, the turbidity remained below the values obtained with the unmodified LCD, presumably because of the large size of the droplets and their settling to the bottom of the well. Consistent with the formation of protein-dense condensates, sedimentation assays showed an enhanced accumulation of both unmodified TDP-43^LCD^ and SUMO2-TDP-43^LCD^ in the pellet fraction, while SUMO1-TDP43^LCD^ largely remained in the supernatant (**Fig. 1d, S1e**).

The surface wetting behaviour of the SUMO2 fusion protein suggested a highly liquid state of the condensates^32,33^. Indeed, FRAP assays of SUMO2-TDP-43^LCD^ confirmed an enhanced liquidity compared to the unmodified LCD (**Fig. 1e**). Nevertheless, in both cases, condensates slowly solidified over time, judging from the observed loss of mobility. To assess whether SUMO would affect the propensity of TDP-43^LCD^ to aggregate, we employed a fluorescent dye, Proteostat, which labels aggregates via non-specific intercalation^34^, and induced *in vitro* aggregation by spiking in 1% of pre-aggregated protein. While fusion of SUMO1 had little effect on the aggregation of TDP-43^LCD^ over time, SUMO2 completely abolished aggregate formation over the course of the experiment (**Fig. 1f**), suggesting that the mobilising effect of SUMO2 on the LCD might protect condensates from transition to solid aggregates.

Even at substoichiometric ratios, the SUMO isoforms had marked effects on TDP-43^LCD^ phase separation (**Fig. S1f**), indicating that they can still modulate condensate formation when not all molecules in the population carry the modifier.

The striking differences between the effects of SUMO1 and SUMO2 on TDP-43^LCD^ were surprising, given the similarity of the SUMO isoforms in terms of structure and biological function. To probe their origin, we conducted coexistence simulations with the fusion proteins using an amino acid-level coarse-grained (CG) model, CALVADOS2^35–37^. This model represents peptides with one bead per amino acid, mapped on the Cα atoms, and enables the tracing of how macroscopic changes, such as the partitioning into protein-dense and protein-dilute phases, arise from residue-level interactions. Simulations were conducted with 100 protein chains at 300K. A representative snapshot of the simulation box (**Fig. 1g**) shows that unmodified TDP-43^LCD^ forms a stable, condensed aggregate. Under the same conditions, the pre-formed SUMO1-TDP-43^LCD^ system lacked stability and systematically dissolved over the simulation timescale, with individual molecules detaching from the dense phase and disseminating into the surrounding dilute phase. In contrast, the SUMO2-TDP-43^LCD^ system developed a larger and denser condensed phase. To quantify these findings, we calculated the saturation concentrations (*C*_*sat*_), representing the protein concentration in the dilute phase that is in equilibrium with the dense phase. Consistent with the opposing effects of the isoforms, we observed an increase in *C*_*sat*_ for SUMO1-TDP-43^LCD^ and a decrease for SUMO2-TDP-43^LCD^ relative to the unmodified TDP-43^LCD^ (**Fig. S1g-h**).

Taken together, these data indicate that SUMO1 strongly inhibits phase separation of TDP-43’s LCD, whereas SUMO2 promotes condensate formation and at the same time prevents aggregation by enhancing the liquidity of the condensates. Importantly, both effects require the covalent attachment of the modifier to the LCD.

### SUMO1 and SUMO2 have opposite effects on TDP-43^LCD^ in cells

Compared to the defined conditions of the *in vitro* assays, the cellular environment is crowded and much more complex. To determine the effects of the SUMO isoforms on TDP-43 in this milieu, we expressed TDP-43^LCD^ and the SUMO fusions with a C-terminal eGFP-tag at high levels via transient transfection (**Fig. 2a**), ensuring comparable protein levels by western blotting (**Fig. S2a**). We quantified the fraction of cells forming condensates (n > 3 foci per cell) as well as the average number of foci per cell and their average size (**Fig. 2b-e**). Under these conditions, TDP-43^LCD^ alone exhibited a moderate degree of condensate formation. Consistent with the results of the *in vitro* assays and the MD simulations, SUMO1 effectively prevented droplet formation, whereas SUMO2 strongly enhanced phase separation according to all criteria. Solubility assays with total cell lysates showed that both SUMO1 and SUMO2 prevented the partitioning of TDP-43^LCD^ to the detergent-insoluble pellet (**Fig 2f-g, S2b**).

**Fig 2:**
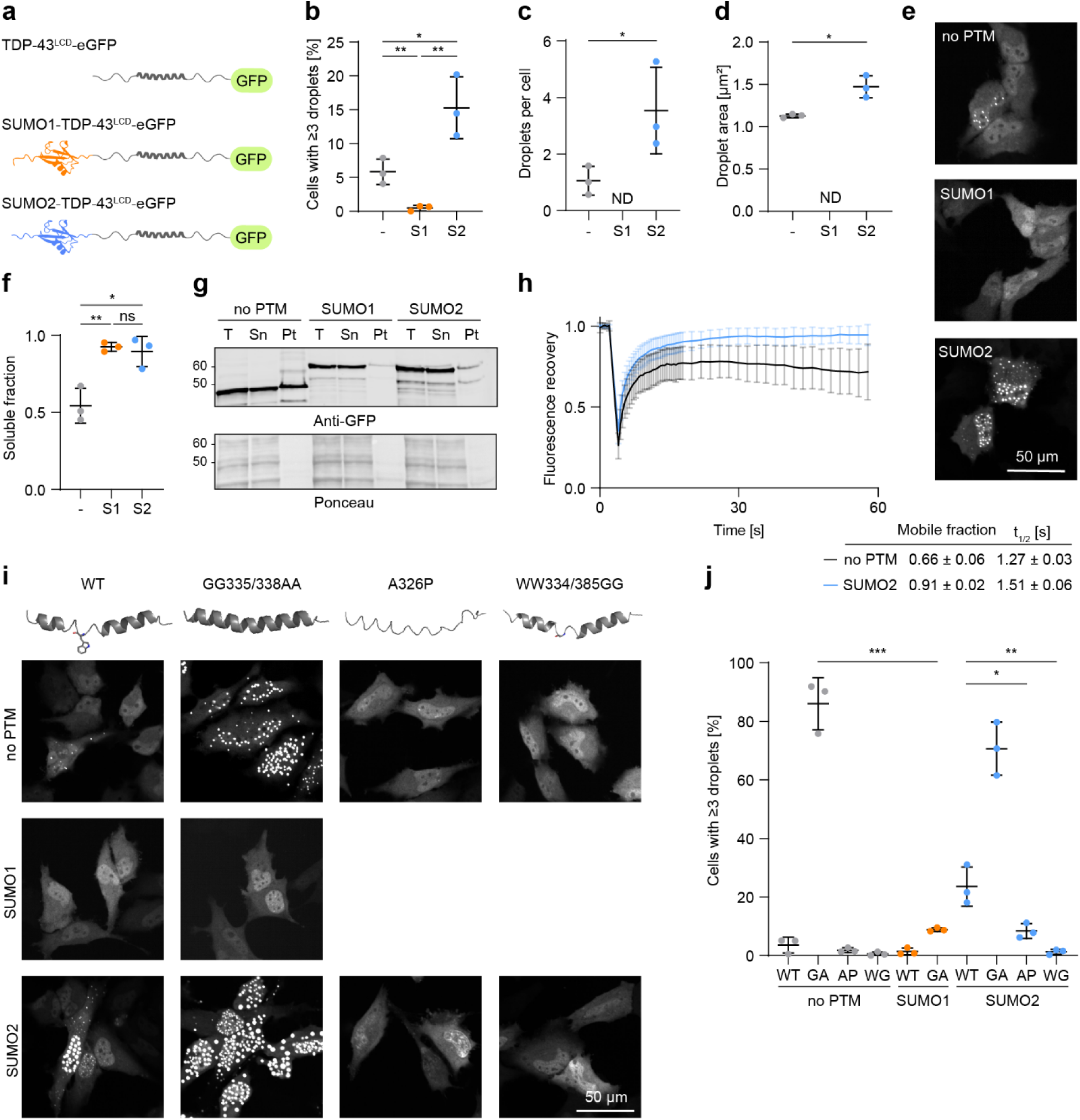
Effects of SUMO on TDP-43^LCD^ in cells resembles those *in vitro*. **a,** Schematics of TDP-43 constructs used for assays in cells. **b-e,** SUMO1 inhibits condensate formation of TDP-43’s LCD in cells, while SUMO2 enhances it. Quantification of droplet formation monitored by live-cell fluorescence microscopy of HeLa cells transfected with TDP-43^LCD^-eGFP fused to SUMO isoforms. Values represent the means ±SD of three independent replicates with at least 1,000 imaged cells per experiment (two-tailed paired Student’s t-test, *: *p* < 0.05, **: *p* < 0.005; ND: not determined). **b**, Fractions of transfected cells forming n ≥ 3 foci. **c**, Average number of droplets per cell. **d**, Average droplet size. **e,** Representative images. **f-g,** Both SUMO1 and SUMO2 solubilise TDP-43^LCD^ in cells. Sedimentation assays with lysates of HeLa cells transfected with TDP-43^LCD^-eGFP constructs, analysed by SDS-PAGE and western blotting. **f,** Quantification of band intensities as a measure of solubility (Sn/(Sn+Pt)), showing means ±SD of three independent experiments. **g,** representative blot (T: total protein before centrifugation; Sn: supernatant; Pt: pellet after centrifugation). Ponceau staining served as loading control. See Fig. S2b for additional replicates. **h**, SUMO2 mobilises TDP-43^LCD^. Exemplary FRAP assays of HeLa cells transfected with TDP-43^LCD^-eGFP constructs. Values represent means ±SD of approximately 10 bleached droplets in one experiment. Means ±SD of mobile fractions and half-life (t_1/2_) values from three independent experiments are shown in the table. **i-j,** SUMO isoforms respond to the helical propensity TDP-43^LCD^. Expression controls are shown in Fig. S2c. **i,** AlphaFold 3 models of the TDP-43^LCD^ helical region with the indicated mutations (top). Representative live-cell fluorescence microscopy images of HeLa cells transfected with TDP-43^LCD^-eGFP phase-separation mutants (bottom). **j**, Fractions of transfected cells forming n ≥ 3 foci. Values represent the means ±SD of three independent replicates with at least 1,000 imaged cells (two-tailed paired Student’s t-test, *: *p* < 0.05, **: *p* < 0.005, ***: *p* < 0.0005).

When we analysed the mobility within droplets by FRAP assays, we observed that SUMO2 strongly enhanced liquidity, resulting in a mobile fraction of 91% compared to 66% for the unmodified LCD (**Fig. 2h**). This implies that SUMO2 can render TDP-43^LCD^ condensates more liquid-like even in the crowded environment of the cell.

To assess the strength with which the SUMO isoforms impinge on TDP-43^LCD^ phase separation, we introduced a set of mutations with defined effects on condensate formation. TDP-43’s LCD contains a conserved, partially helical region (CR, amino acids 320-343, **Fig. 1a**) that is critical for self-interaction, thus promoting phase separation but also modulating aggregation^38^. Accordingly, mutants with an enhanced helical propensity, such as GG335/338AA, strengthen phase separation^39^, while helix-disrupting mutations, such as A326P, have a negative effect^38^. In addition to the helical propensity, a pair of tryptophane residues (W334, W385) promotes self-interaction and phase separation^40^. When we compared droplet formation of the SUMO-TDP-43^LCD^ constructs, we found that the enhanced helical propensity of the GG335/338AA mutation promoted phase separation of both isolated TDP-43^LCD^ and the SUMO2 fusion, whereas SUMO1 strongly attenuated this effect (**Fig. 2i-j, S2c**). SUMO2 in turn was able to partially compensate for the helix-breaking effect of the S326P mutation but did not restore droplet formation of the WW334/385GG double mutant. This implies that the positive effect of SUMO2 relies at least to some degree on the self-interaction of the helical region in the LCD.

In summary, our observations suggest a strong negative effect of SUMO1 on the phase separation of TDP-43^LCD^ even under conditions where self-interaction is facilitated. In contrast, SUMO2 promotes phase separation as long as the LCD maintains a certain level of self-interaction.

### SUMO1 inhibits TDP-43^LCD^ phase separation via intramolecular contacts

To elucidate the structural basis of SUMO1’s inhibitory action on TDP-43^LCD^ phase separation, we turned to hydrogen-deuterium exchange mass spectrometry (HDX-MS). Considering its strong effect on the helix-enhancing LCD mutant (**Fig. 2i**), we hypothesized that SUMO1 might physically block self-assembly of the LCD and that this might result in altered solvent accessibility in regions critical for the contact. We therefore measured the differential deuterium uptake by the SUMO1-TDP-43^LCD^-MBP fusion protein relative to that of an equimolar mixture of free SUMO1 and TDP-43^LCD^-MBP. In these assays, the MBP unit was left attached to prevent phase separation of the unmodified LCD. Peptide coverage exceeded 90% for both SUMO1 and TDP-43^LCD^ (**Fig. S3a-c**). Compared to the mixture of the free proteins, the fusion protein exhibited several distinct changes in differential deuterium uptake, suggestive of altered solvent accessibility or hydrogen bonding (**Fig. 3a, S3a-c**).

**Fig 3:**
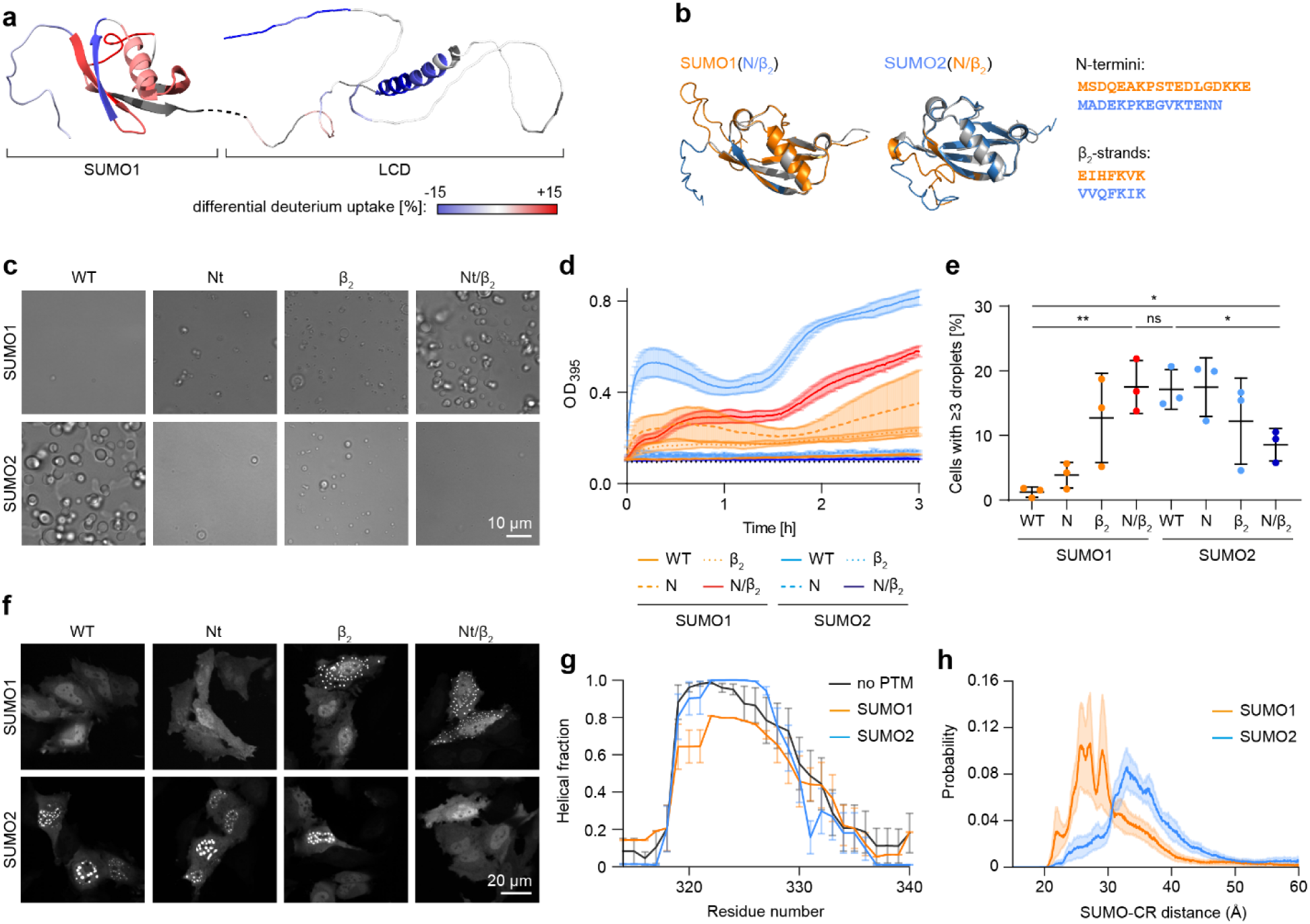
SUMO1 prevents TDP-43^LCD^ phase separation by blocking its self-assembly sites. **a,** Fusion of SUMO1 to TDP-43^LCD^ causes regional changes in solvent accessibility and secondary structure (hydrogen bonding) in both proteins. Differential deuterium uptake by SUMO1-TDP-43^LCD^-MBP compared to a non-covalent mixture of SUMO1 and TDP-43^LCD^-MBP after 30 s of deuterium exposure, measured by HDX-MS and plotted onto AlphaFold 3 models of the two parts. Residues not identified by MS are shown in dark grey. **b,** Swapping of the N-termini and β_2_-strands between SUMO1 and SUMO2 does not grossly interfere with structure. AlphaFold 3 models of SUMO hybrids (grey with colour-coded critical regions), superimposed onto structural models of the respective isoforms (coloured). Sequences are shown on the right. **c,** SUMO’s effects on TDP-43^LCD^ phase separation are driven by its N-terminus and β_2_-strand. Condensate formation of TDP-43^LCD^ (50 µM) fused to SUMO hybrids as indicated, monitored by DIC microscopy 1 h after addition of TEV protease. TEV cleavage patterns are shown in Fig. S3d. **d,** Turbidity measurements of TDP-43^LCD^ (50 µM) fused to SUMO hybrids as indicated, after addition of TEV protease. Values represent means ±SD of three technical replicates (one of three independent experiments). **e-f,** Effects of SUMO2 hybrids are reproduced in cells. **e,** Fractions of HeLa cells expressing TDP-43^LCD^-eGFP fused to SUMO hybrids forming n ≥ 3 condensates. Values represent the means of three independent replicates with at least 1,000 imaged cells (two-tailed paired Student’s t-test, * *p*<0.05; ** *p*<0.005; ns: not significant). **f,** Representative live-cell fluorescence microscopy images. **g,** Fusion of SUMO1 to TDP-43^LCD^ decreases the helical propensity of its self-interacting region. Atomistic MD simulations of TDP-43^LCD^ and its SUMO fusions, showing the helical fraction of the self-interacting region (CR, aa 314-340). For each construct, 10 independent trajectories of 1 μs were run, and the final 800 ns of each replica were used for analysis. **h,** SUMO1-TDP-43^LCD^ adopts a more compact conformation than the SUMO2 fusion. Probability distribution of the distance between the centre of mass of the SUMO moiety and that of the self-interacting region (CR) from the same atomistic MD trajectories as in panel g.

Within the LCD, a decrease in differential deuterium uptake of the helical CR after a 30 s exposure indicated protection of the region responsible for phase separation. In addition, a short segment of the C-terminus, including residue W412, also reported to modulate phase separation^40^, exhibited enhanced protection. Within the SUMO1 moiety, protected regions were located at the flexible N-terminus as well as the second strand of SUMO’s β-sheet (β_2_). This was contrasted by an increased deuterium uptake of the adjacent β1-strand and the opposing α-helix, suggesting allosteric effects and subtle opening of the SUMO1 core. At prolonged exposure times, differences in solvent accessibility largely faded, arguing against any persistent structural alterations (**Fig. S3a-c**).

Our results suggested that SUMO1 contacts the CR within TDP-43’s LCD in an intramolecular fashion via its N-terminus and β_2_-strand, thus preventing the helix-helix interaction that mediates phase separation of the LCD. To verify this, we exchanged the relevant regions of SUMO1 for those of SUMO2. According to AlphaFold 3 modelling, the swap did not grossly affect SUMO’s overall structure (**Fig. 3b**). Consistent with our model, transplanting SUMO2’s N-terminus and β_2_-strand onto the SUMO1 framework enhanced *in vitro* droplet formation in an additive manner, while a SUMO2 chimera containing the corresponding regions of SUMO1 efficiently prevented phase separation in microscopy and turbidity assays (**Fig. 3c-d, S3d**). A similar behaviour was observable in cells (**Fig.3e-f, S3e**).

To elucidate the underlying mechanism at atomistic level, we performed all-atom MD simulations. These indicated a decrease in the helical propensity of the self-interacting CR within TDP-43’s LCD when fused to SUMO1 as compared to the unmodified LCD or the SUMO2 fusion (**Fig. 3g**). This suggests that SUMO1 may influence phase separation not only by sterically occluding the CR interface, but also by perturbing its secondary structure and thereby weakening its ability to act as a structured self-association element. Consistent with the proposed intramolecular engagement of SUMO1 with the CR, distance analyses from the same atomistic trajectories showed lower average separation between the centre of mass of the SUMO moiety and that of the CR for SUMO1-TDP-43^LCD^ compared to SUMO2-TDP-43^LCD^ (**Fig. 3h**). The shift of the distance distribution towards lower values is in line with more frequent and persistent intramolecular contacts in the SUMO1 fusion and is also evident from snapshots of the structures, where SUMO1, but not SUMO2, tended to remain near the CR while simultaneously reducing its helical propensity (**Fig. S3f**).

Overall, these data illustrate how SUMO1, by contacting the CR of TDP-43’s LCD and possibly interfering with its helicity, prevents phase separation of the domain in an intramolecular fashion. SUMO1’s N-terminus as well as its β_2_-strand are critical for this isoform-specific, intramolecular action.

### SUMO2 promotes TDP43^LCD^ phase separation via intermolecular self-interaction

We were unable to acquire HDX-MS data to probe solvent accessibility in the phase-separated state of SUMO2-TDP-43^LCD^. We therefore employed crosslinking mass spectrometry (XL-MS) to capture potential contact sites. To differentiate inter- from intramolecular interactions, we used an equimolar mixture of native (^14^N) and ^15^N-labelled SUMO2-TDP-43^LCD^-MBP^41^. In a first step, a bifunctional crosslinker, Sulfo-SDA, was mono-linked to the proteins via NHS chemistry at varying molar ratios in solution. Phase separation was initiated by treatment with TEV protease for 2 h, and the highly non-selective diazirine group of the crosslinker was activated by UV-A irradiation (**Fig. S4a**). As a negative control, TEV protease was omitted from one set of reactions to prevent phase separation. Analysis by SDS-gel electrophoresis and western blotting revealed distinct new species in the TEV-treated samples (**Fig. 4a**), one of them corresponding in size to the crosslinked dimer of cleaved SUMO2-TDP-43^LCD^ (ca. 60 kDa). Tryptic digestion of this band excised from a Coomassie-stained gel (**Fig. S4b**), followed by mass spectrometric analysis, revealed a large number of crosslinks originating from the SUMO2 moiety with high scores (**Fig. 4b**).

**Fig 4:**
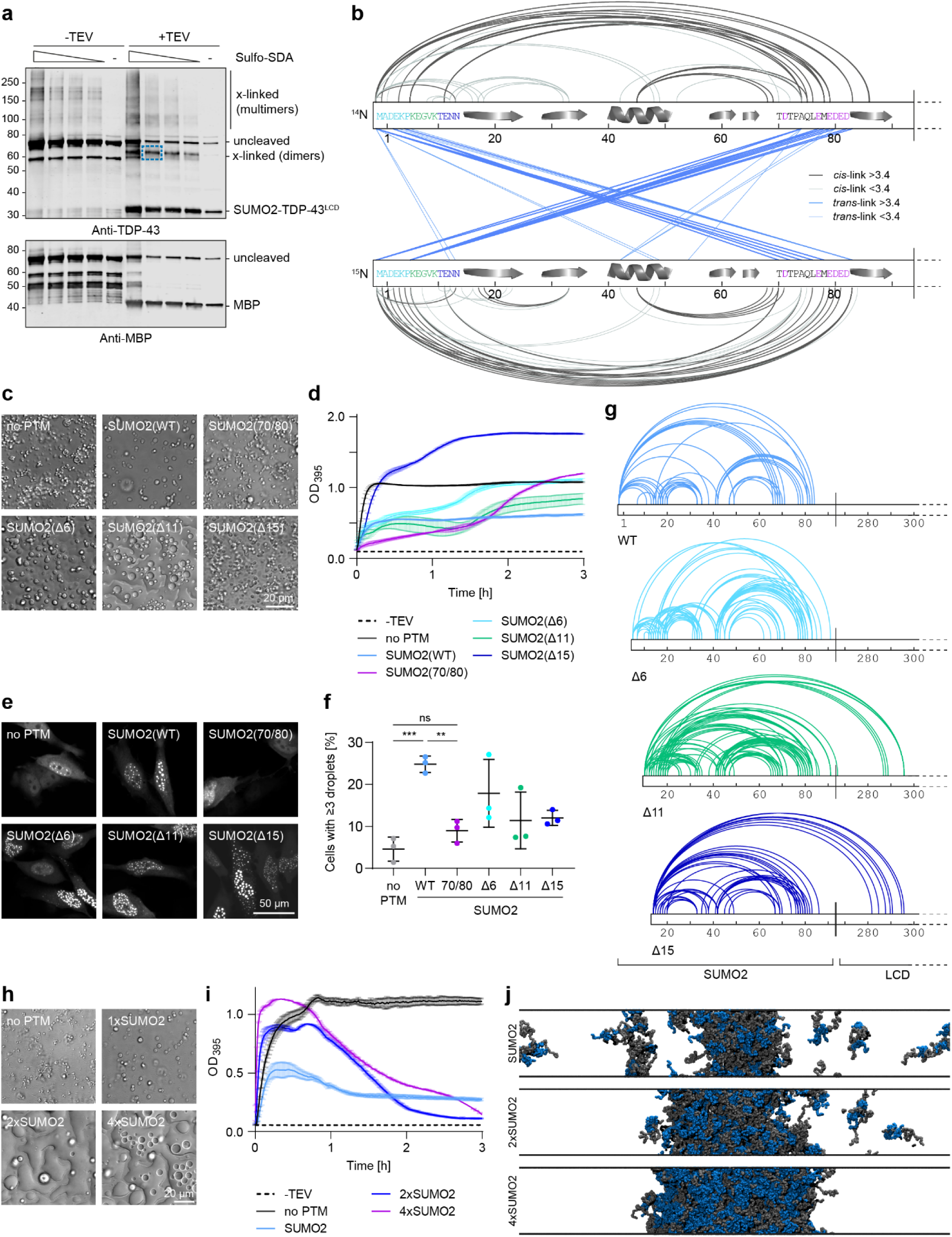
SUMO2 promotes TDP-43^LCD^ phase separation by *trans*-interactions between its N-terminus and the 70/80 region. **a,** Crosslinking of SUMO2-TDP43^LCD^ depends on phase separation. Western blot analysis of Sulfo-SDA crosslinking reactions of ^14^N- and ^15^N-SUMO2-TDP-43^LCD^, applying varying concentrations of the crosslinker. The band used for XL-MS analysis is indicated by a blue box. **b,** Inter- and intramolecular SUMO2 interactions involve defined regions in its N-terminus and 70/80 region. *Cis*- and *trans*-crosslinks plotted according to confidence score onto SUMO2. Secondary structure elements and amino acid sequences of regions identified with high confidence are highlighted. **c,** SUMO2’s 70/80 region is critical for enhancing phase separation, while its N-terminus plays a complex role. Condensate formation of TDP-43^LCD^ (50 µM) fused to the indicated SUMO2 mutants, monitored by DIC microscopy 1 h after addition of TEV protease (70/80 mutant: D71/80/82N, E77/79/81Q). **d,** Turbidity measurements of TDP-43^LCD^ (50 µM) fused to the indicated SUMO2 mutants after addition of TEV protease. Values represent means ±SD of 3 technical replicates in one out of three measurements. TEV protein cleavage patterns are shown in Fig. S4f. **e-f,** Effects of SUMO2 mutants are reproduced in cells. **e,** Representative live-cell fluorescence microscopy images of HeLa cells expressing TDP-43^LCD^-eGFP fused to the indicated SUMO mutants. **f,** Fractions of cells forming n ≥ 3 condensates. Values represent the means ±SD of three independent replicates with at least 1,000 imaged cells (two-tailed paired Student’s t-test, * *p* < 0.05, ** *p* < 0.005). **g,** Truncation of SUMO2’s N-terminus induces novel contacts between the remaining N-terminal tail and TDP-43’s LCD. Sulfo-SDA-induced crosslinks of SUMO2-TDP-43^LCD^ carrying the indicated N-terminal truncations, plotted onto SUMO2 and the N-terminal region of TDP-43^LCD^. Amino acids are numbered according to the original proteins. **h-j**, Enhancement of phase separation scales with the number of SUMO2 moieties. **h,** Condensate formation of TDP-43^LCD^ (50 µM) fused to one, two or four units of SUMO2, monitored by DIC microscopy 1 h after addition of TEV protease. Internal SUMO2 units in the head-to-tail fusion carry the Δ11 truncation. **i,** Turbidity measurements of TDP-43^LCD^ (50 µM) fused to one, two or four units of SUMO2 after addition of TEV protease. Values represent means ±SD of 3 technical replicates in one out of three measurements. TEV cleavage patterns are shown in Fig. S4i. **j**, Representative snapshots from CG slab coexistence simulations with the CALVADOS2 model at 320 K for 100-chain systems of TDP-43^LCD^ fused to one, two or four SUMO2 units, showing progressively thicker and more densely packed condensates with increasing SUMO2 valency.

Among these, the most prominent signals were found between the flexible N-terminus and a loop connecting the fourth and fifth strand of SUMO’s β-sheet, previously dubbed the “70/80” region^42^. Intriguingly, this interaction was detectable both *in cis* and *in trans*, suggesting intra- as well as intermolecular interactions between the two sites. In contrast, interactions within or between the LCDs were hardly detectable, most likely due to the scarcity of lysine residues in this domain and consequently the lack of efficient NHS-mediated attachment of the Sulfo-SDA crosslinker in the first step of the crosslinking (**Fig. S4c**). Analysis of a higher-order crosslinked species of ca. 120 kDa yielded a similar pattern (**Fig. S4b,d**).

To verify the relevance of the identified SUMO2 regions for enhancing phase separation, we introduced a set of 6 mutations neutralising the negative charge of the 70/80 region (D71/80/82N, E77/79/81Q), and we constructed a series of N-terminal truncations (Δ6, Δ11, Δ15) (**Fig. S4e**). Microscopy (**Fig. 4c**) and turbidity measurements (**Fig. 4d, S4f**) confirmed that the 70/80 mutation abolished the effect of SUMO2 on TDP-43^LCD^, bringing droplet sizes down to the level of the unmodified LCD. The effects of the N-terminal truncations were more difficult to interpret. Deletion of the first 6 or 11 amino acids enhanced droplet sizes, with Δ11 forming extremely large, liquid-like patches, but deletion of the entire unstructured N-terminus (Δ15) resulted in amorphous structures that generated extremely high turbidity (**Fig. 4c-d**). In cells, the 70/80 mutation had similar effects as *in vitro*, whereas all N-terminal truncations enhanced droplet formation relative to unmodified TDP43^LCD^ (**Fig. 4e-f, S4g**). XL-MS analysis of the N-terminal truncation mutants Δ11 and Δ15 now indicated a series of contacts between the new N-terminus of the SUMO moiety and the LCD, but outside the helical region (**Fig. 4g, S4h**). These new interactions may explain the complex effects of the shortened N-termini.

Our results suggested that SUMO2 contributes to phase separation by providing additional contact sites for multivalent interactions. Based on this model, increasing the degree of SUMOylation should even further enhance its effect on condensate formation. Indeed, expanding the number of N-terminal SUMO2 moieties to two or four via head-to-tail fusions strongly accelerated phase separation and enhanced droplet size and liquid-like behaviour of the SUMO2_n_-TDP-43^LCD^ protein *in vitro* (**Fig. 4h-i, S4i**). For these experiments, we used the Δ11 mutant for the internal moieties to mimic the geometry of natural SUMO2 chains, which are predominantly linked via K11^43,44^. CG coexistence MD simulations using CALVADOS2 confirmed the propensity of di- or tetra-SUMO2 fusions to TDP-43^LCD^ to form more densely packed condensates, which protein chains being progressively recruited into a single dense phase and the surrounding dilute phase becoming visibly depleted of protein (**Fig. 4j**). Consistent with these observations, the corresponding slab density profiles revealed a systematic decrease in the saturation concentration (*C_sat_*) with increasing SUMO2 valency (**Fig. S4j**). Note that simulations were performed at 320 K rather than the 300 K used in the earlier CALVADOS2 simulations to weaken overall attractive interactions and thereby raise the *C_sat_* of the mono-SUMO2 fusion, allowing changes in phase behaviour upon increasing SUMO2 valency to be quantified more accurately.

Taken together, these data indicate that SUMO2 undergoes self-interactions, mediated by contacts between its flexible N-terminus and its 70/80-loop, that can contribute to phase separation by enhancing multivalency. Interestingly, the degree of this effect appears to be modulated by a balance between inter- and intramolecular interactions involving the two critical regions.

### Effects of SUMO1 and SUMO2 are reproduced with full-length TDP-43

To assess the physiological relevance of our results, we asked whether the effects of the SUMO isoforms on the LCD would also apply to full-length TDP-43. Moreover, TDP-43 is known to undergo proteolytic cleavage, resulting in C-terminal fragments of ca 35 and 25 kDa (CTF35 and CTF25) that are thought to be particularly aggregation-prone and have been implicated in disease^45,46^. We therefore fused the SUMO isoforms to the N-termini of full-length TDP-43, CTF35 and CTF25 and expressed them as GFP-tagged proteins in HeLa cells to quantify droplet formation (**Fig. 5a, S5a**). In this assay, SUMO1 had a similar inhibitory effect on all constructs as on the isolated LCD (**Fig. 5b**). In contrast, SUMO2 did not enhance the number of droplets as expected; in fact, it reduced droplet formation of full-length TDP-43, possibly due to the dominating effect of TDP-43’s N-terminal oligomerisation domain^47^. Likewise, in sedimentation assays, SUMO2’s influence on solubility was limited, and a significant solubilisation was only observed with CTF25, while SUMO1 enhanced the solubility of all three constructs to a similar degree as for the LCD (**Fig. 5c-d**). Nevertheless, FRAP assays showed that both SUMO isoforms enhanced the mobility of full-length TDP-43 condensates and decreased the recovery time (t_1/2_), with SUMO1 having a slightly stronger effect than SUMO2 (**Fig. 5e**).

**Fig 5:**
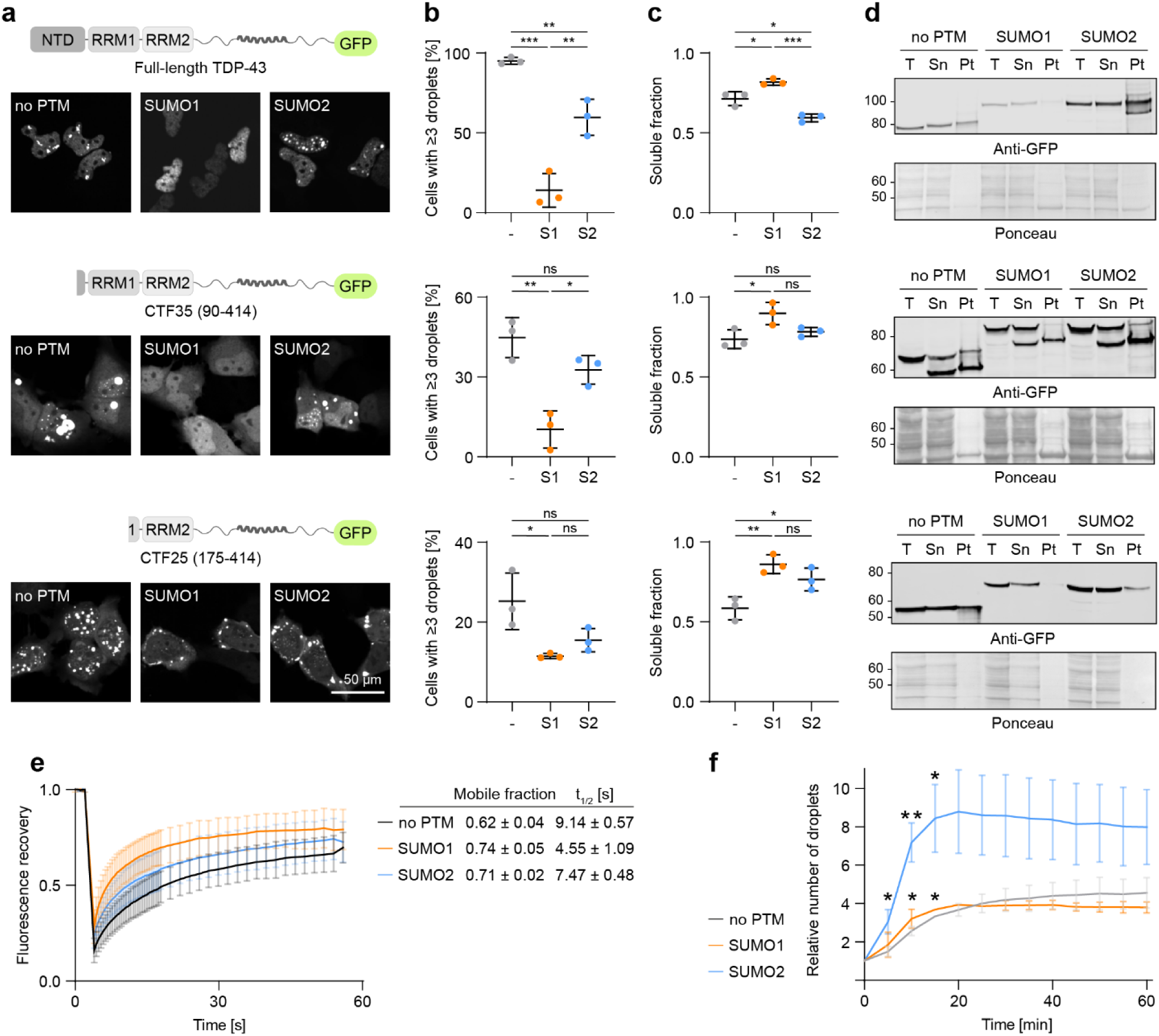
Effects of SUMO isoforms on phase separation in cells apply to full-length TDP-43 and its C-terminal fragments. **a,** Representative live-cell fluorescence microscopy images of HeLa cells transfected with full-length TDP-43-eGFP (top) or its C-terminal fragments, CTF35-eGFP (middle) and CTF25-eGFP (bottom), fused to SUMO isoforms. **b,** Fractions of HeLa cells transfected with the constructs shown in panel a forming n ≥ 3 condensates. Values represent the means ±SD of three independent replicates with at least 1,000 imaged cells (two-tailed paired Student’s t-test, * p < 0.05, ** p < 0.005, *** p < 0.0005). **c-d,** Sedimentation assays with lysates of HeLa cells transfected with the constructs shown in panel a, analysed by SDS-PAGE and western blotting. **c,** Quantification of band intensities as a measure of solubility (Sn/(Sn+Pt)) from three independent experiments (means ±SD). **d,** Representative blots. Ponceau staining served as loading control. **e,** FRAP assays of HeLa cells transfected with full-length TDP-43-eGFP fused to SUMO isoforms. Values represent means ±SD of approximately 30 bleached droplets from three independent biological replicates. Means ±SD of mobile fractions and t_1/2_ values are shown in a table. **f,** SUMO2 enhances stress-induced TDP-43 condensation in cells. Quantification of nuclear bodies of TDP-43-eGFP fused to SUMO isoforms in HeLa cells (average numbers ±SD relative to the unstressed condition (t=0) from three independent experiments) after exposure to oxidative stress (100 µM NaAs_2_) (One-way ANOVA with Tukey’s multiple comparison, * *p* < 0.05, ** *p* < 0.005).

Building on these results, we set out to examine the effects of SUMO in a more physiological setting, where TDP-43 condensate formation is driven by cellular stress. After exposure to oxidative stress by treatment with sodium arsenite (NaAsO_2_), TDP-43 reversibly assembles in dynamic, phase-separated nuclear bodies^48^. Consistent with its mobilising effect, fusion of SUMO2 drastically accelerated the formation of these structures and also their numbers relative to the unstressed condition (**Fig. 5f, S5b**). SUMO1 had a milder effect and did not affect the relative number of nuclear bodies, as it decreased their absolute numbers to similar degrees in the absence or presence of stress (**Fig. S5c**).

In summary, SUMO1 reduced condensate formation of all TDP-43 variants tested, highlighting the importance of its blocking effect on the self-association of the helical region within the LCD. While SUMO2 exhibited variable effects on the degree of condensate formation, its mobilising effect within condensates was reproducible for full-length TDP-43.

### SUMO2 enhances phase separation of various IDRs

The mechanisms by which the two SUMO isoforms impinge on TDP-43 suggested that SUMO1 acts very specifically on the LCD, while the effect of SUMO2 should be largely independent of the protein that it is attached to. To test these assumptions, we used the same TEV-cleavable MBP fusion strategy as before to construct SUMO fusions to three other IDRs known to undergo SUMOylation and phase separation, but with distinct characteristics (**Fig. 6a**). A protein corresponding to exon 1 of Huntingtin (HttEx1) is found enriched in pathological inclusions of Huntington’s disease patients^49^. It is composed of an N-terminal helical domain (N17) and a semi-helical structure of glutamine (Q) residues, interrupted by a proline-rich stretch^50^, and undergoes phase separation and aggregation in a manner depending on the number of Q residues, with a critical threshold of 40 Q for a disease phenotype^49^. HttEx1^Q23^, chosen for our assays, forms condensates *in vitro* in the presence of a crowding agent, 20% dextran, at 20 µM^51^. Another disease-relevant IDR-containing factor is the RNA-binding protein FUS, which – like TDP-43 – has been associated with ALS and FTD^52–54^. One of its IDRs, FUS^QGSY^, which is enriched in polar amino acids and prone to form fibrils^55^, shares prion-like characteristics with TDP-43^LCD^ but apparently forms condensates without adopting a transient secondary structure^56^. A second IDR, FUS^RGG1^, cooperates with a folded RRM domain in nucleic acid binding^57^ and uses its “RGG” tripeptide motifs for phase separation via the stickers-and-spacers principle^58^. For each of these IDRs, we adjusted the concentrations of the protein and a crowding agent, PEG-8,000, to achieve a moderate degree of phase separation in turbidity assays and by DIC microscopy (**Fig. 6b-c**, **S6a**). As a negative control, we mixed the IDRs with equimolar amounts of free SUMO, which – as expected – did not affect phase separation (**Fig. S6b-d**). In line with our model postulating a general effect of SUMO2, covalent fusion of this isoform enhanced condensation of all three IDRs, producing large, round droplets (**Fig. 6b-c**).

**Fig 6:**
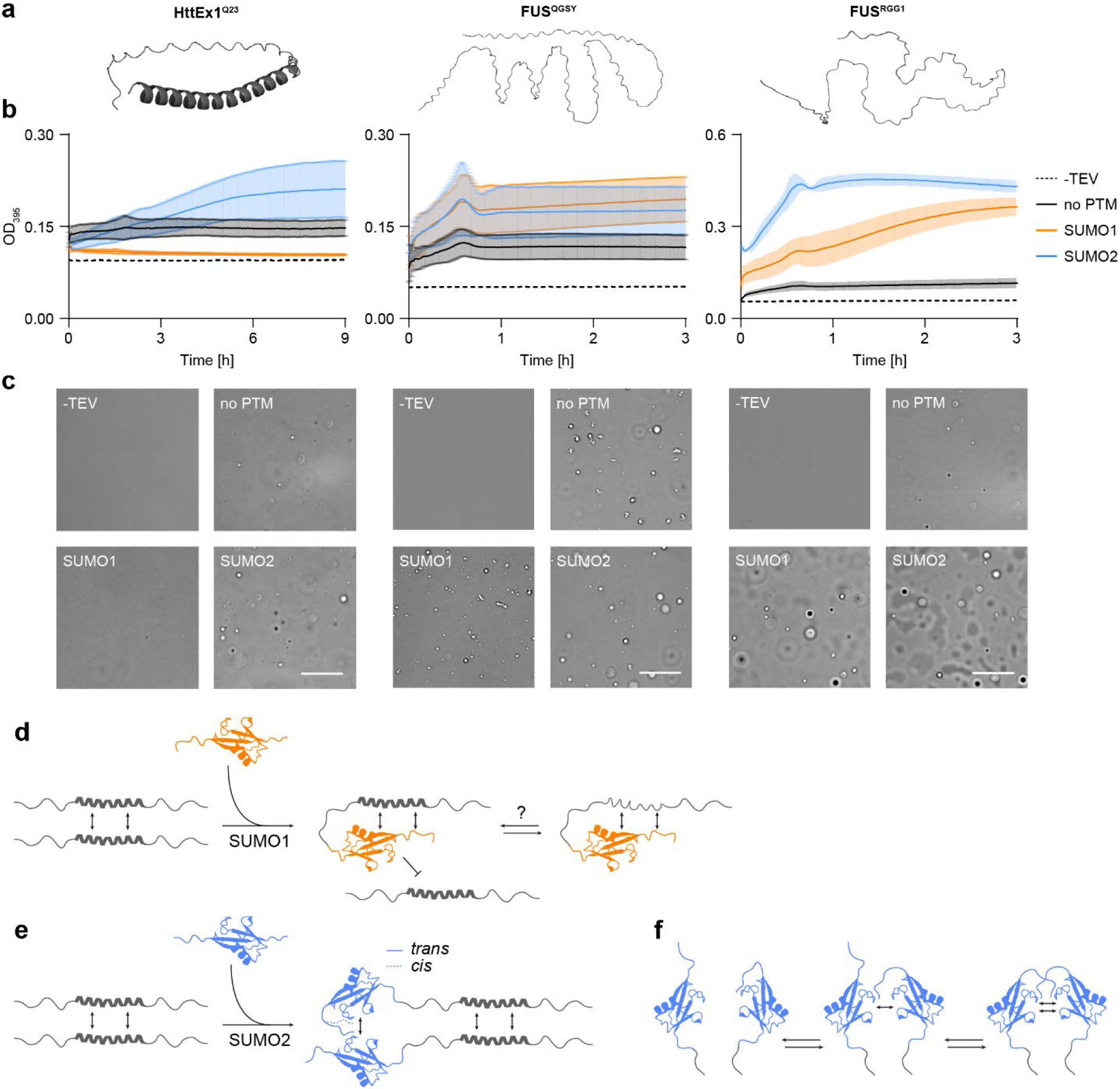
SUMO2 promotes phase separation of various different IDRs. **a,** AlphaFold 3 models of Huntingtin Exon1 (HttEx1^Q23^, aa 1-89) and the IDRs of FUS (FUS^QGSY^, aa 1-163; FUS^RGG1^, aa 164-285). **b,** Turbidity measurements of the indicated domains fused to SUMO isoforms after addition of TEV protease (HttEx1^Q23^: 20 µM in 10% (w/v) PEG-8,000; FUS^QGSY^ and FUS^RGG1^: 10 µM in 5% (w/v) PEG-8,000). Values represent means ±SD of 3 technical replicates (one out of three measurements). TEV cleavage patterns are shown in Fig. S6a. **c,** DIC microscopy of HttEx1^Q23^, FUS^QGSY^, and FUS^RGG1^ under similar conditions as in panel b (time points: HttEx1^Q23^, 6 h; FUS^QGSY^ and FUS^RGG1^, 1 h). **d,** Proposed model for the intrinsic effects of SUMO1 on TDP-43. **e,** Proposed model for the intrinsic effects of SUMO2 on phase-separating IDRs. Note that for the sake of clarity, the interaction between SUMO2’s N-terminus and the 70/80-loop has been illustrated unidirectionally, although a mutual interaction between two SUMO2 moieties is conceivable. **f,** Proposed model for the competition between inter- and intramolecular interactions involving SUMO2’s N-termini and 70/80-loops.

Surprisingly, however, SUMO1 exerted opposite effects on HttEx1^Q23^ and the FUS-derived IDRs (**Fig. 6b-c**): while HttEx1^Q23^ was prevented from condensation in a manner reminiscent of TDP-43^LCD^, the effect of SUMO1 on FUS^QGSY^ and FUS^RGG1^ was more comparable to that of SUMO2 in that phase separation was enhanced. Interestingly, the two FUS IDRs were affected differently by the modifiers, with FUS^RGG 1^ forming much larger and rounder condensates than FUS^QGSY^, where particularly SUMO1 led to more irregularly shaped droplets.

In summary, our results support our hypothesis that the enhancement of IDR phase separation by SUMO2 is independent of the nature of the conjugated protein, while the effect of SUMO1 is more specific to the properties of its target.

## Discussion

SUMOylation is well known to modulate protein phase separation and aggregation in cells. In this study, we show that these activities go far beyond the well-characterised SUMO-SIM interactions, relying on inter- and intramolecular contacts between SUMO’s flexible N-terminus and defined regions within its structurally conserved core. Our data suggest a mechanistic basis for the effects of the two SUMO isoforms, SUMO1 and SUMO2, on a model IDR, TDP-43^LCD^, and other unstructured domains.

### SUMO isoforms modulate phase separation and aggregation via intrinsic effects

SUMO1 exerts a strongly inhibitory effect on TDP-43^LCD^ phase separation *in vitro* and in cells, mediated by intramolecular contacts of a region comprising its N-terminus and its β_2_-strand with the helical region of the LCD responsible for self-association (**Fig. 6d**). These interactions appear to prevent the self-association that drives phase separation and aggregation of the LCD. This may be mediated by a physical blocking of the helix-helix interaction and/or by a full or partial disruption of the helical structure. The notion that helix-stabilising mutations (GG335/338AA) were not effective in restoring phase separation upon the SUMO1-TDP-43^LCD^ fusion argues against a strong influence of SUMO1 on helical structure, whereas MD simulations implied a partial loss of helicity in this construct.

Although we did not pinpoint the effect of SUMO1 to specific residues within its N-terminus and β_2_-strand, comparison with the respective regions of SUMO2 reveals a stronger net negative charge within the N-terminus and a pair of exposed polar or charged residues (E33 and H35 *versus* V29 and Q31) in SUMO1, which could mediate the contacts to the helical region of the LCD. Testing mutations within the LCD itself was not feasible as it would have been difficult to separate their effects on SUMO1 interaction from their consequences on helicity and self-association. Of note, despite the inhibitory effect of SUMO1 on phase separation, fusion of this isoform did not prevent aggregation of TDP-43^LCD^ *in vitro*. This confirms previous data demonstrating LCD fibrillation in conditions incompatible with phase separation^59^ and suggests that phase separation is not an obligatory intermediate on the path to aggregation. Our findings indicate that the SUMO1-TDP-43 interaction cannot be generalised to other IDRs but appears to be specific for TDP-43. Interestingly, however, SUMO1 also inhibited phase separation of HttEx1^Q23^, which – like TDP-43 – contains a helical region. Whether the underlying mechanism is the same and SUMO1 preferentially interacts with α-helices, remains to be determined.

In contrast to SUMO1, SUMO2 promotes phase separation of all IDRs tested, producing large droplets and rendering condensates more mobile and liquid-like. Our data indicate that this is driven by intermolecular contacts between SUMO2’s N-terminus and a set of negatively charged residues in the 70/80 region of a second molecule of SUMO2 (**Fig. 6e**). We suggest that these interactions enhance the multivalency of any IDR to which SUMO2 is attached. Corresponding intramolecular contacts between the N-terminus and the 70/80-loop had been reported for SUMO2 before, although their significance was not discussed^42^. Thus, competition between inter- and intramolecular interactions (**Fig. 6f**) may render the SUMO2-SUMO2 interaction extremely dynamic. Such competition might also underlie the behaviour of the N-terminal truncation mutants, where *trans*-interactions might still be possible while *cis*-interactions may be compromised, which would favour the intermolecular contacts. We propose that the dynamic nature of these interactions contributes to the general solubilising effect of SUMO2.

### Dynamic contacts involving the N-termini define SUMO’s activities

Despite the ambiguous behaviour of SUMO1 on its fusion partners, the two isoforms share several features relevant to their respective activities. First, the interactions driving their effects on the IDRs are extremely weak, as they only become apparent when the modifiers are covalently attached to their targets. This stands in contrast to the previously described non-covalent inhibition of α-synuclein aggregation by SUMO1, mediated by a SIM-dependent interaction in the low micromolar range^60^. Second, the disordered N-termini of both SUMO isoforms apparently do not adopt random positions but exhibit a tendency to contact distinct sites within the SUMO body in an intramolecular fashion. While SUMO2’s N-terminus prefers the acidic 70/80 region, SUMO1’s N-terminus has been reported to align with the SIM-binding groove, thereby interfering with SUMO-SIM interactions^42^. Interestingly, removal of negative charge from SUMO1’s N-terminus shifts its preference towards the 70/80 region^42^. Thus, it is conceivable that the N-terminus of SUMO1 might also engage in intermolecular SUMO1-SUMO1 interactions. In this case, the positive effect of SUMO1 on phase separation of the FUS-derived IDRs may follow a similar principle of enhancing multivalency as we found for SUMO2, although the relevant contact region on the SUMO body might differ.

SUMO’s N-termini have previously been implicated in solubilising SUMOylated proteins by forming a disordered “cloud” around them, thus preventing aggregation-prone interactions by following a principle of “entropic bristles”^61^. Our data confirm the importance of the flexible N-termini but argue for the contributions of more defined interactions involving the SUMO core.

### Both SUMO isoforms prevent aggregation of TDP-43 in cells

A multitude of studies have identified TDP-43 as a target of both SUMO1 and SUMO2^22,23,25–29^. Yet, the effects of the different isoforms have not been compared. Two SUMO-specific E3s, PIAS1 and PIAS4, have been implicated in TDP-43 modification in association with nuclear PML bodies and cytoplasmic stress granules, respectively^26,27^. However, neither of the E3s has been reported to differentiate between the SUMO isoforms. This raises the question of whether the opposing effects of SUMO1 *versus* SUMO2 on TDP-43 phase separation are physiologically meaningful. A possible answer follows from the notion that global SUMO2/3 conjugation is predominantly induced by cellular stress, whereas SUMO1 is thought to represent more basal conditions^13,62,63^. Thus, modification of TDP-43 with SUMO1 would contribute to protection against aggregation in unstressed cells by reducing or preventing its phase separation. Upon induction of the stress response, when TDP-43 undergoes phase separation and associates with stress granules, enhanced modification with SUMO2 would prevent the transition to aggregates by rendering the resulting condensates more liquid-like. Moreover, the enhanced mobility of SUMO2-TDP-43 might accelerate its compartmentalisation into PML bodies, which would in turn facilitate its RNF4- and TOPORS-mediated turnover in this compartment^26^. Overall, a protective effect of both SUMO paralogues on TDP-43 is therefore conceivable, although it appears to be accomplished by different mechanisms.

### Outlook

Our results underscore the importance of SUMO’s intrinsic properties for phase transitions of TDP-43 and other targets. However, additional factors, such as SUMO’s “classical” SIM-dependent mode of interaction, its subcellular localisation, and the presence of other posttranslational modifications, including phosphorylation and acetylation, clearly contribute their own effects on phase separation and solubility. Elucidating their complex interplay will be necessary to fully understand the extent to which the biophysical characteristics of SUMO impinge on its targets. Moreover, quantifying SUMOylation in an isoform-specific manner without perturbing its endogenous balance is still challenging but will be important to monitor TDP-43 SUMOylation at its physiological modification sites under basal and stressed conditions in cells. Finally, although many intrinsically disordered proteins linked to neurodegenerative diseases, including TDP-43, FUS, Huntingtin, Tau and α-synuclein, are targets of SUMOylation, there is still controversy about whether the modification is generally beneficial or toxic^14^. Despite these challenges, our study opens potential avenues for harnessing SUMO’s protective activity on TDP-43 and other disordered proteins for pharmacological interventions, for example by inhibiting relevant deSUMOylating enzymes or via the design of small molecules that modulate SUMO’s properties.

## Methods

### Plasmid construction

Plasmids were generated using restriction cloning, Gibson Assembly or site-directed mutagenesis. All plasmids were propagated in *E. coli* TOP10 and verified by Sanger sequencing. A list of plasmids used in this study is provided in Table S1.

For soluble overexpression of IDRs in *E. coli*, a modified pET vector was generated. A 3C-protease-cleavable His_6_-FKBP and a TEV-protease-cleavable MBP-tag were positioned N-and C-terminally, respectively, from the IDR coding sequence for purification purposes and for maintaining solubility, respectively. For TDP-43, a single cysteine (G275C) was introduced by site-directed mutagenesis as described before^11^. SUMO1 or SUMO2 were cloned as linear fusions between the 3C-site and the IDR. A mutation of SUMO’s C-terminal glycine (G) to valine (V) was introduced to enhance stability. For production of untagged SUMO1 and SUMO2, the coding sequences were cloned into a pCoofy1 vector^64^.

For transient overexpression of TDP-43 derivatives in mammalian cells, a modified pEGFP vector was generated, and relevant portions of TDP-43 were inserted between a constitutive CMV promoter and a C-terminal eGFP moiety. SUMO1 or SUMO2 were cloned as linear fusions upstream of the TDP-43 coding sequences, including the C-terminal G-to-V mutation.

### Production of recombinant proteins

All IDPs were expressed in *E. coli* BL21 DE3 CodonPlus (DE3)-RIPL (Agilent). Cultures were grown LB medium at 37 °C to an OD_600_ of ca. 0.6, briefly cooled down, and expression was induced with 0.5 mM IPTG at 16 °C overnight. For production of ^15^N-labelled SUMO2-TDP-43^LCD^-MBP, cells were grown in M9-minimal medium [4 g/l glucose, 0.5 g/l ^15^NH_4_Cl, trace elements (100x, Sigma), 100 µM CaCl_2_, 1 mM MgSO_4_, 22 mM KH_2_PO_4_, 42 mM Na_2_HPO_4_, 86 mM NaCl] and induced as stated above. Cells were pelleted and lysed in IMAC buffer (30 mM Tris pH 8, 300 mM NaCl, 15 mM imidazole) supplemented with 1 mM DTT, 5 mM MgCl_2_, Protease Inhibitor (Roche), Sm nuclease (in-house) by passing them through a pressure homogeniser system (Constant Systems) at 1.8 kbar. The lysate was supplemented with 0.1% (v/v) Triton X-100, cleared by centrifugation, and incubated with Ni-NTA agarose (Qiagen) for 2 h at 4 °C. After washing 5x with IMAC buffer, beads were incubated with 3C-protease (in-house) for 1 h at room temperature (RT), removing His_6_-FKBP and releasing the MBP-tagged IDPs from the column. The flowthrough was concentrated and subjected to gel filtration on a Superdex200 16/600 SEC column (Cytiva) equilibrated in storage buffer (20 mM HEPES pH 7.4, 150 mM NaCl, 10% (v/v) glycerol, 1 mM DTT), followed by concentration to 100 µM. Proteins were snap-frozen in liquid nitrogen and stored at -70 °C.

For fluorescent labelling of TDP-43^LCD^ constructs, single-cysteine proteins were purified as stated above, but using 0.5 mM TCEP instead of 1 mM DTT in the storage buffer. Samples were incubated with a 2-fold molar excess of AzDye488 for 2 h at room temperature (RT), followed by quenching with 2 mM DTT. Excess dye was removed by another round of gel filtration in storage buffer. The degree of labelling was determined using a spectrophotometer. For fluorescence microscopy and FRAP analysis, 5%-labelled protein samples were generated by mixing TDP-43^LCD^ and labelled TDP-43^LCD(G275C)^ using freshly thawed stocks.

SUMO proteins were expressed in *E. coli* Rosetta 2(DE3)pLysS. The purification procedure is similar as stated for IDRs. After lysis by pressure homogenisation and IMAC, 3C-protease (in-house) was used for removing the His_6_-tag and releasing untagged SUMO with a N-terminal GP-remnant from the column. The flowthrough was subjected to gel filtration on a Superdex75 16/600 pg column (Cytiva), followed by concentration to 200 µM and snap-freezing.

### Protein analysis by Coomassie staining and western blotting

Protein samples were separated using 4-12% Bis-Tris gels. Proteins were either visualised by Coomassie staining or transferred onto a nitrocellulose membrane for western blotting. Blots were blocked for 1 h in 5% skim milk. The following primary antibodies were used: Anti-GFP (1:1,000, Roche, #11814460001), Anti-MBP (1:10,000, NEB, #E8032S), Anti-TDP-43 (1:1,000, Proteintech, #12892-1-AP). After incubation overnight at 4 °C, blots were briefly washed and incubated with fluorophore-tagged secondary antibodies (1:10,000; IRDye® 800CW donkey anti-mouse, IRDye® 800CW goat anti-rabbit IgG, IRDye® 680RD donkey anti-mouse; LI-COR) for 1 h at RT with three subsequent washes. Blots were imaged using a LI-COR Odyssey fluorescence scanner system.

### Human cell culture

HeLa cells were grown in DMEM (Gibco) supplemented with 10% (v/v) FBS, 2 mM L-glutamine, 100 U/ml penicillin, and 100 µg/ml streptomycin at 37 °C, 5% CO_2_ and 85% relative humidity. Transient transfection was performed using FuGENE (Promega) according to the manufacturer’s instructions. Protein expression was allowed for at least 24 h before further processing. For oxidative stress treatment, cells were exposed to 0.1 mM NaAsO_2_ for up to 1 h.

### Preparation of human cell lysates

For protein analysis in total lysates, cells from a 6-well plate (Corning) were harvested and lysed in RIPA buffer (HEPES pH 7.4, 100 mM NaCl, 2.5 mM MgCl_2_, 0.1% (w/v) SDS, 0.5% (w/v) Sodium deoxycholate, 1% (w/v) NP-40) supplemented with protease inhibitor cocktail (Roche) and Sm nuclease (in-house). After incubation for 15 min at 4 °C, protein concentrations were determined by BCA assay (Thermo Fisher Scientific), samples were normalised and incubated in 4x LDS buffer (Invitrogen) at 95 °C for 5 min.

### Live-cell imaging

Transiently transfected cells on a 96-well PhenoPlate (Revvity) were imaged using an Opera Phenix high-throughput spinning-disc confocal microscope (Perkin Elmer) equipped with a 40x water objective with autofocus in confocal mode. For visualisation of nucleus and cytoplasm, cells were stained with DRAQ5 (Invitrogen) at a final concentration of 5 µM. eGFP signals were imaged with 100% laser power for 40 ms, and DRAQ5 with 100% laser power for 100 ms. For each tested condition, at least 1,000 cells were imaged at three planes.

Condensate formation was quantified with an automated pipeline using the built-in evaluation software. Cell regions (membrane, nuclei, cytoplasm) were detected using the DRAQ5 signal. eGFP intensity was calculated, and positively transfected cells were selected if the median intensity value was above 125. Spots were separately detected in nuclei and cytoplasm. The number of cells harbouring more than 3 droplets were divided by the total number of transfected cells as a measure of phase separation. Statistical analysis was performed in GraphPad Prism 8.3.0.

### DIC and fluorescence microscopy of purified proteins

Purified proteins were diluted to 10-50 μM in an appropriate phase separation buffer at a final volume of 50 μl. The reaction mix was transferred into 18-well μ-Slides with glass bottom (Ibidi), and phase separation was induced by adding 0.1 mg/ml TEV-protease for 50 µM IDR. For lower IDR concentrations, TEV amounts were appropriately scaled. Droplets were imaged after 1 h at RT using an AF7000 widefield microscope (Leica) with a 63x/1.4 oil immersion objective and TL-DIC. For monitoring phase separation by fluorescence microscopy, 5%-labelled protein mixtures were used as described above. This preparation was imaged using an L5 filter cube (Ex: 460-500, Em: 512-542).

### Turbidity assays

Purified proteins were diluted to 10-50 μM in an appropriate phase separation buffer at a final volume of 20 μl. Reaction mixes were transferred into black polystyrene clear bottom 384-well microplates (Corning). Phase separation was induced by the addition of 0.1 mg/ml TEV-protease for 50 μM IDR (or proportionally lower amounts for lower IDR concentrations). Plates were sealed with PCR foil to avoid evaporation. Turbidity was assessed by measuring the OD_395_ using a Spark M20 microplate reader (Tecan) at 25 °C for 1-3 h. The Z-position was determined directly from a reference well containing 20 µl of phase separation buffer.

### Aggregation assays

To enhance the propensity of protein aggregation, phase separation of TDP-43^LCD^-MBP was induced as stated above, followed by shaking at 37 °C and 1,400 rpm for 3 days to generate aggregated protein. In 20 µl reactions, 50 µM of TDP-43^LCD^-MBP was mixed with 1% pre-aggregated protein, and Proteostat (Enzo) reaction components were added following the manufacturer’s instructions. Plates were sealed with PCR foil to avoid evaporation. Proteostat fluorescence was assessed by measuring emission at 600 nm after excitation at 550 nm using a SpectraMax iD5 plate reader and low gain settings at 37 °C for 24 h.

### FRAP assays

Fluorescence recovery after photobleaching was performed using a Stellaris 8 FALCON laser scanning confocal microscope (Leica) with an integrated FRAP module, using a 63x oil immersion objective. Images were recorded at a resolution of 256x256 pixels, a scanning rate of 600 Hz, and a pinhole size of 1 AU. Raw data were analysed using EasyFRAP^65^.

For measuring FRAP *in vitro*, phase separation of 5% fluorescently labelled protein was induced as stated above for fluorescence microscopy assays. A white light laser set to 480 nm at 0.4% power was used for imaging, and a 488 nm laser at 10% power was used for photobleaching. Bleaching regions were selected to cover approximately 50% of a droplet. Images were recorded every 0.5 s, with three images pre-bleaching, ten images bleaching and 240 images post-bleaching. Additional ROIs were selected after imaging to cover the entire droplet and its surrounding region (background).

For measuring FRAP in cells, transiently transfected cells were exposed as stated above, but with 30 images taken every 0.5 s post-bleaching, followed by 20 images taken every 2 s to reduce phototoxicity. Additional ROIs were selected to cover the entire cell and its background (inter-cell space).

### Sedimentation and solubility assays

Sedimentation assays were performed as described before^66^. Briefly, for *in vitro* sedimentation assays monitoring the degree of phase separation, condensation was induced as stated above for turbidity measurements, and protein samples were incubated for 1 h at 25 °C. Reactions were centrifuged for 15 min at 21,000x g and 4 °C. The supernatant was mixed with 4x LDS buffer (Invitrogen), while the pellet was resuspended in 1x LDS buffer. For sedimentation assays from cell material, which are used to assess protein solubility rather than phase separation, total lysates were prepared from transiently transfected cells in RIPA buffer as stated above. An input sample (5%, T) was collected, and the remainder of the lysate was centrifuged for 15 min at 21,000x g and 4 °C. The supernatant (Sn) was removed, and pellets (Pt) were washed and resuspended in denaturing buffer (30 mM Tris-HCl pH 8.5, 7 M urea, 2 M thiourea, 4% CHAPS). Fractionated samples were mixed with 4x LDS buffer. Statistical analysis was done in GraphPad Prism 8.3.0.

### Hydrogen-deuterium exchange mass spectrometry – sample preparation

Hydrogen-deuterium Exchange mass spectrometry (HDX-MS) was performed for the detection of binding interfaces and conformational changes upon SUMO1 fusion to TDP-43^LCD^ in comparison to free SUMO1 mixed with TDP-43^LCD^. For these assays, the MBP unit was left attached to prevent phase separation of the unmodified LCD. Equilibration (E)-, labelling (L) and quench (Q)-buffer were prepared in either H_2_O (buffer E: 20 mM HEPES, 150 mM NaCl, pH 7.5; buffer Q: 150 mM potassium phosphate, pH 2.2) or D_2_O (buffer L: 20 mM HEPES, 150 mM NaCl, pH 7.1). Prior to the experiment, buffer Q was cooled to 0 °C and buffers E/L were equilibrated to RT. SUMO1-TDP-43^LCD^-MBP and TDP-43^LCD^-MBP were adjusted to a final concentration of 15 µM, corresponding to 25 pmol on the protease column per injection for HDX experiments with and without addition of free SUMO1 at equimolar concentration. TDP-43^LCD^-MBP was preincubated with free SUMO1 for 30 min at RT.

### HDX-MS – data acquisition

HDX-MS experiments were performed using an automated HDX-2 system (Waters, Milford USA) as described previously^67^. HDX reactions were initiated by incubation of 4 µl SUMO1-TDP43^LCD^-MBP (or TDP-43^LCD^-MBP + free SUMO1) with 56 µl of either buffer E (reference, 0 s) or buffer L (D_2_O labelling) for multiple time points (30, 120, 600 and 2,100 s). The exchange reactions were quenched by mixing 50 µl of the sample with 50 µl of buffer Q. After 30 s of incubation at 0 °C, 95 µl were injected into a temperature-controlled chromatography system equipped with a 50 µl sample loop (HDX nanoAqcuity UPLC, Waters). Protein digestion was conducted using a nepenthesin-2/pepsin column (AffiPro, AP-PC-006).

Following protein digestion, peptides were trapped on a C18 pre–column (C18 1.7 µM VanGuard 2.1 x 5 mm pre-column; Waters) at a flow rate of 100 µl/min for 3 min and separated using an analytical reversed-phase column (C18 1.7 µM Acquity UPLC 1 x 100 mm reverse phased column; Waters) with a 7-min linear gradient from 5% acetonitrile (ACN) to 40% ACN + 0.23% formic acid (FA) at 40 µl/min. Next, the ACN concentration was rapidly increased to 95%, maintained for 2 min, prior to column (re-)equilibration with 95% H_2_O + 0.23% FA for 2 min. The reversed-phase chromatography system was cooled to 0 °C to minimise HD back-exchange. Peptides were measured using a Synapt G2-Si mass spectrometer (Waters, Milford USA) in HDMSE mode (50-2,000 m/z) on the basis of retention time, mass-to-charge ratio and precursor ion mobility (LC, m/z, IM). The mass spectrometer was fitted with an electrospray ionisation source equipped with an additional independent LockSpray probe (GluFib lock mass: 785.8426 m/z) for alternating lock mass infusion.

### HDX-MS – data analysis and statistics

Peptide identification was performed using the ProteinLynx Global Server 3.0.3. (PLGS, Waters) in both non-deuterated reference conditions (SUMO1-TDP-43^LCD^-MBP fusion and TDP-43^LCD^-MBP + free SUMO1) with separate peptide searches for comparative downstream analysis of TDP-43^LCD^-MBP and SUMO1, respectively. Peptides with a confidence score >6, a length of 3 to 20 amino acids, and an RSD of retention time below 10%, that were identified in at least six out of eight technical replicates were considered for evaluation.

Annotation of all isotopic clusters for each peptide was performed using HDExaminer (Sierra Analytics, Trajan Automation). Relative deuterium uptake was calculated through comparison of centroid masses of deuterated isotope clusters of each peptide and the corresponding non-deuterated reference. All acquired spectra were manually inspected and strictly revised as necessary, excluding the MBP-tag from analysis. For the SUMO1-TDP-43^LCD^-MBP fusion condition, one technical replicate per labelling timepoint (30, 120, 600 and 2,100 s) was excluded based on data structure and experimental reproducibility.

Peptides that exhibit statistically significant differences in deuterium uptake (Student’s t-test with a 95% confidence interval, p≤ 0.05) in at least 2 out of 4 (TDP-43^LCD^) or 3 out of 4 (SUMO1) timepoints were used for evaluation and visual presentation on the structural models for TDP-43^LCD^ and SUMO1 using UCSF Chimera^68^.

### Crosslinking mass spectrometry – sample preparation

Crosslinking mass spectrometry (XL-MS) was performed for the detection of *cis*- and *trans*-interactions within and between SUMO2-TDP-43^LCD^ in the phase-separated state. The hetero-bifunctional crosslinker Sulfo-SDA (Thermo Fisher Scientific), containing a 3.9 spacer arm, was used in all cases. SUMO2-TDP-43^LCD^-MBP was mixed with ^15^N-labelled SUMO2-TDP-43^LCD^-MBP at equimolar ratio to achieve a total protein concentration of 50 µM in storage buffer after addition of the crosslinker. For a simplified experiment with the N-terminally truncated SUMO2-TDP-43^LCD^ mutants, only unlabelled (^14^N) proteins were used. Sulfo-SDA was prepared in serial dilutions to cover molar crosslinker:protein ratios between 50:1 and 1:1 in storage buffer. Protein and crosslinker solutions were mixed and incubated at RT for 45 min to enable the reaction of the NHS function group. Without further quenching, 0.1 mg/ml TEV-protease was added, and the reaction was further incubated for 2 h. The Diazirine-group was activated by exposing the reaction mixes to 5,000 mJ/cm^2^ UV-A. The samples were subsequently mixed with 4xLDS buffer and separated by SDS-PAGE.

After Coomassie staining and identification of crosslinked bands, excised gel pieces were cut into small cubes, followed by destaining in 50% ethanol and 25 mM ammonium bicarbonate. Proteins were then reduced in 10 mM DTT at 56 °C and alkylated by 50 mM iodoacetamide in the dark at RT. Subsequently, proteins were digested by the addition of 1 µg trypsin per sample in 50 mM ammonium bicarbonate at 37 °C overnight. Following peptide extraction sequentially using 30% and 100% ACN, the sample volume was reduced in a centrifugal evaporator to remove residual ACN. Following acidification by FA, the resultant peptide solution was purified by solid phase extraction in C18 StageTips^69^.

### XL-MS – data acquisition, ^14^N-/^15^N-SUMO2 interactions

Peptides were separated via an in-house packed 45-cm analytical column (inner diameter: 75 μm; ReproSil-Pur 120 C18-AQ 1.9-μm silica particles, Dr. Maisch GmbH) on a Vanquish Neo UHPLC system (Thermo Fisher Scientific). The online reversed-phase chromatography separation was conducted through a 70-min non-linear gradient of 1.6-32% ACN in 0.1% FA at a nanoflow rate of 300 nl/min. Eluted peptides were sprayed directly by electrospray ionisation into an Orbitrap Astral mass spectrometer (Thermo Fisher Scientific).

Mass spectrometry was conducted in data-dependent acquisition mode using a top50 method with one full scan in the Orbitrap analyser (scan range: 325 to 1,300 m/z; resolution: 120,000, target value: 3 × 10^6^, maximum injection time: 25 ms) followed by 50 fragment scans in the Astral analyser via higher energy collision dissociation (HCD; normalised collision energy: 26%, scan range: 150 to 2,000 m/z, target value: 1 × 10^4^, maximum injection time: 10 ms, isolation window: 1.4 m/z). Precursor ions of unassigned, +1 or higher than +6 charge state were rejected. Additionally, precursor ions already isolated for fragmentation were dynamically excluded for 15 s.

### XL-MS – data acquisition, ^14^N-SUMO2 variant interactions

Peptides were analysed using an Orbitrap Exploris 480 mass spectrometer (Thermo Fisher Scientific) coupled to a FAIMS Pro Interface and a Vanquish Neo UHPLC system. Peptides were separated by online reversed phase chromatography through a 90-min gradient of 1.6-32% ACN with 0.1% FA at a nanoflow rate of 300 nl/min. The eluted peptides were sprayed directly by electrospray ionisation into the mass spectrometer via the FAIMS device. Each sample was measured twice using different compensation voltage (CV) settings. Mass spectrometry was conducted in data-dependent acquisition mode through a combination of internal and external CV steppings (-45, -50, -55, -60, -65, -70 V). A top10 method was applied with one full scan (resolution: 60,000, scan range: 325-1,300 m/z, target value: 3 × 10^6^, maximum injection time: 50 ms) followed by 10 fragment scans via higher energy collision dissociation (HCD in stepped mode^70^: normalised collision energy 22, 26, 30%; resolution: 30,000, scan range: 150-2,000 m/z, target value: 1 × 10^5^, maximum injection time: 50 ms, isolation window: 1.4 m/z). Only precursor ions of +3 to +6 charge state were selected for fragment scans. Additionally, precursor ions already isolated for fragmentation were dynamically excluded for 20 s.

### XL-MS data analysis, ^14^N-/^15^N-SUMO2 interactions

Peak lists as mzML files were extracted from the raw data using ProteoWizard^71^. An MS2 denoise filter was applied to pick 20 most abundant peaks in a moving window size of 100 Da. The peak lists were then searched using the SIM-XL software (version 1.5.7.2)^41^. The crosslinker specificity for Sulfo-SDA (mass shift: +82.041865 Da) was assigned as linking any amino acid to K, S, T, Y residue or the protein N-terminus. Mass tolerance was 5 ppm for precursor ions and 10 ppm for fragment ions. Trypsin/P and full enzyme specificity were assigned. Up to 3 missed cleavages were allowed. Carbamidomethylation at cysteine residue was chosen as fix modification. Methionine oxidation was considered as variable modification. An FDR threshold was set to 5% for the target-decoy search. Homodimer crosslink analysis specifically for the non-labelled and ^15^N-labelled versions of SUMO2-TDP-43^LCD^ was performed considering 2 isotopic possibilities. MBP-derived peptides were included in this analysis. *Cis*- and *trans*-links were visualised by xiNET^72^.

### XL-MS data analysis, ^14^N-SUMO2 variant interactions

Mass spectra from individual CV settings were extracted^73^. Spectra were then pre-processed, and the peak lists were extracted in MaxQuant (version 2.1.3.0) as previously described^74^, ignoring a de-noising filter. The peak lists (*.HCD.FTMS.sil0.apl files) were searched using xiSEARCH (version 1.8.7)^75^ against a target-decoy database consisting of the protein sequence of the corresponding recombinant protein and TEV protease. Trypsin/P specificity was assigned. Up to 4 missed cleavages were allowed. Crosslink search included SDA (mass shift: + 82.0418648147 Da) linking any amino acid to K, S, T, Y residue or the protein N-terminus and noncovalently associated peptides^76^. The MS-cleavability of the diazirine-based crosslinker was also taken in account^77^. Carbamidomethyl on cysteine was assigned as fixed modification. Variable modifications included methionine oxidation, loop link and mono-link with an unreacted or hydrolysed end. Mass tolerance was 5 ppm for MS1 and 6 ppm for MS2. Residue-level FDR was set to 1%.

### AlphaFold 3 simulations

Peptide sequences were submitted to the AlphaFold 3 Server with seeding set to Auto^78^. Models were visualised using PyMOL 3.1.

### Molecular dynamics simulations

To dissect the physicochemical principles governing biomolecular condensates, we used a dual-resolution molecular dynamics (MD) framework comprising: (i) fully atomistic explicit-solvent simulations and (ii) residue-level CG simulations with the CALVADOS2 model^35–37^. The CALVADOS2 level identified sequence-encoded drivers of phase separation, which then guided atomistic simulations to resolve the inter- and intramolecular interaction networks underpinning emergent mesoscale organisation. The description of each model and simulation methodology is described below.

#### Atomistic Simulation

Initial coordinates for the post-translational modifiers were obtained from the Protein Data Bank (PDB) (SUMO1: 2N1V; SUMO2: 2N1W). The structure of TDP-43^LCD^ (aa 267–414) was sourced from the AlphaFold Protein Structure Database (AF-Q13148-F1). Each chain was first simulated using the CG model described in the next section. Then, we used 10 structures from single-chain CG simulation of 5 *μ*s and back-mapped them to atomistic structures using the PULCHRA module^79^. Each protein system was subsequently solvated in a dodecahedral box using the TIP4P-D water model^80,81^ with a minimum distance of 2.0 nm between solute atom and the periodic box edge. To approximate physiological ionic strength, the net charge of each protein was first neutralised with counterions, after which NaCl was added to a final concentration of 150 mM. The Amber a99SB-disp force field^80,81^ was employed to describe the interaction and partial charges of the proteins. All the bonds involving hydrogen atoms were constrained with the LINCS algorithm^82^. Temperature was maintained at 300 K using the stochastic velocity rescaling thermostat^83^ with a temperature coupling time constant of *τ*_*T*_ = 0.1 ps. Pressure was controlled at 1 bar using the Parrinello–Rahman barostat^84,85^ with a temperature coupling time constant *τ*_*P*_ = 2 ps. Short-range nonbonded interactions used a 1.0 nm cutoff, while long-range electrostatics were handled using the Particle Mesh Ewald (PME) method^86^. Each system was energy-minimised in two stages consisting of 10^4^ steps of steepest descent followed by 10^4^ steps of conjugate gradient, with a convergence criterion of a maximum force below 10∼*kJ mol*^−1^ *nm*^−1^. The systems were then equilibrated through a multi-stage process. First, a 1 ns constant particles-volume-temperature (NVT) simulation was run with positional restraints (1,000∼*kJ mol*^−1^ *nm*^−2^) on all heavy atoms. This was followed by a 1 ns constant particles-pressure-temperature (NPT) simulation under the same restraint conditions. The force constants were sequentially reduced progressively from 500 to 100, 50, and finally 1∼*kJ mol*^−1^ *nm*^−2^ with each simulation running for 1 ns. Finally, 10 independent production simulations were conducted for 1 *μ*s each, running under the NPT ensemble T = 300 K and P = 1 bar.

#### Coarse-Grained (CG) Simulations

Amino Acid level CG simulations were performed using both HPS^87^, HPS-urry^88^, and CALVADOS2 CG potentials. HPS potential parameters gave an unphysiologically high critical temperature for phase separation with Tc∼370 K. In comparison, CALVADOS2, augmented with cation-pi interactions^89^, reproduced the experimental phase behaviour of the protein with a critical region around 310–320 K, close to physiological conditions, which led us to use the parameter as an optimal choice for the CG simulation. Each amino acid residue was mapped to a single interaction bead interacting through total potential energy, *U*_*tot*_, which decomposed as,

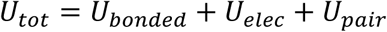

where *U*_*bonded*_ represents bonded connectivity, *U*_*elec*_ encodes screened Coulombic interactions, and *U*_*pair*_ accounts for short-range hydropathy-mediated interactions.

Bonded interactions were described via a harmonic restraint,

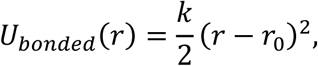

with *k* = 8368 *kJ*. *mol*^−1^. *nm* ^−2^ and *r* _0_ = 0.38 *nm*.

Non-bonded hydrophobic interactions were employed using the Ashbaugh–Hatch potential^90^,

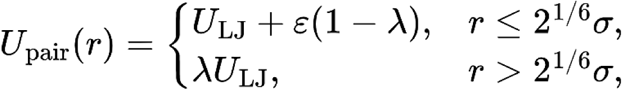

where the *U*_*LJ*_(*r*) is the Lennard-Jones potential given by,

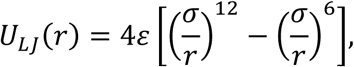

with *ɛ* = 0.8368∼*kJ*. *mol*^−1^, and *λ* determined from sequence-specific parameters. For all simulations in this work, the Ashbaugh–Hatch potential was truncated and shifted at a distance of 2 nm.

Electrostatics were treated using a Debye–Hückel screening model,

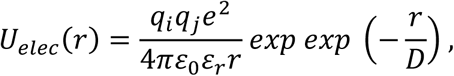

with the Debye length 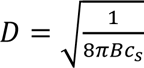 and a temperature-dependent dielectric constant *ɛ* (*T*) given by,

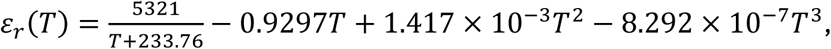

yielding *ɛ*_*r*_(300 *K*) = 77.7. Long-range electrostatic interactions were truncated and shifted at a distance of 4 nm as suggested in the original parametrisation of the CALVADOS2 paper. Since this CG model was parametrised for a disordered protein, we additionally implemented an elastic network to represent the structured region. Each amino acid in the structured region was connected using a harmonic bond with all its neighbouring beads. Root mean squared fluctuation of each amino acid from atomistic simulation was fitted with the elastic network to find the optimal values of cutoff and spring constant. The best fitted Chi-squared value of the elastic network gives a cutoff value of 9 *nm* and an elastic force constant of 600 *kJ*. *mol*^−1^. *nm* ^−2^. We prevent large interaction-induced conformational changes, but this is justified because SUMO1 and SUMO2 adopt a highly conserved, ubiquitin-like fold whose backbone remains essentially unchanged. For the transiently structured CR region in TDP-43^LCD^, we performed simulations both with and without an elastic network. The starting structure of each chain was initialised by linearly conjugating SUMO1(PDB: 2N1V) or SUMO2 (PDB: 2N1W) with the AlphaFold structure of TDP-43^LCD^. The elastic network was applied to residues 22–92 of SUMO1, 18–88 of SUMO2, and residues 319–341 of the TDP-43^LCD^ construct, which were treated as structured. In addition, the elastic network on the CR region of TDP-43^LCD^ was removed in the SUMO1 conjugate, as both atomistic simulations and experimental data indicate that SUMO1 binding disrupts the structural integrity of this region. To map the phase behaviour of the disordered proteins, we utilised a two-phase slab approach in which a dense slab with interfaces perpendicular to the z-axis exchanges material with the surrounding dilute phase. Coexisting densities were then used to determine the binodal and the critical temperature. Initially, 100 protein chains were inserted in a large simulation cell, and the simulation cell was scaled in all direction to ∼20 nm in 500,000 steps. Subsequently, the z-dimension was extended to 280 nm. Production simulations were performed at multiple temperatures (270K-330K) for 5 μs in the NVT ensemble using a Langevin dynamics (ref 18). The integration time step was 10 fs. The first 1 μs was discarded as equilibration as the saturation concentration converged and showed no systematic drift after few hundreds of ns, and the remaining 4 μs were used for analysis. Simulations were carried out with HOOMD-blue v2.9.6.

Phase coexistence densities were extracted by centring the slab configuration along z to maximise,

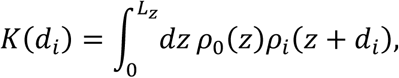

and fitting the averaged density profile *ρ*(*z*) to,

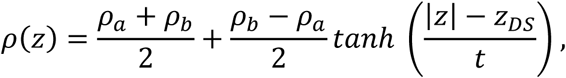

where *ρ*_*a*_ and *ρ*_*b*_ correspond to the protein-rich and dilute phases, respectively.

To obtain the saturation concentration, *C_sat_*, we first computed the z-resolved concentration profile from the coexistence simulations. We then averaged the concentration over all bins of 1 nm assigned to the dilute phase. The reported *C_sat_* is the mean of these dilute-phase bins, and the error bar indicates the standard deviation across the same set of bins.

## Acknowledgements

The authors would like to thank Dorothee Dormann, Saskia Hutten, and Ronald Wong for helpful advice, the Dormann and Beli labs for sharing protocols, plasmids, and reagents, Friedolin Kielisch for advice with statistical analysis, and Ivan Đikić for co-supervision of J.V. We are grateful for support from IMB’s Media Lab and the Core Facilities for Proteomics, Microscopy and Protein Production as well as Projects Z01 and Z02 of CRC 1551. We acknowledge funding from the Deutsche Forschungsgemeinschaft (DFG) for CRC1551 “Polymer concepts in cellular function” (project #464588647) and for the Orbitrap Astral system (project #524805621; Proteomics Core Facility at IMB Mainz).

## Data availability

The mass spectrometry proteomics data have been deposited to the ProteomeXchange Consortium via the PRIDE^91^ partner repository (http://www.ebi.ac.uk/pride) with the following identifiers:

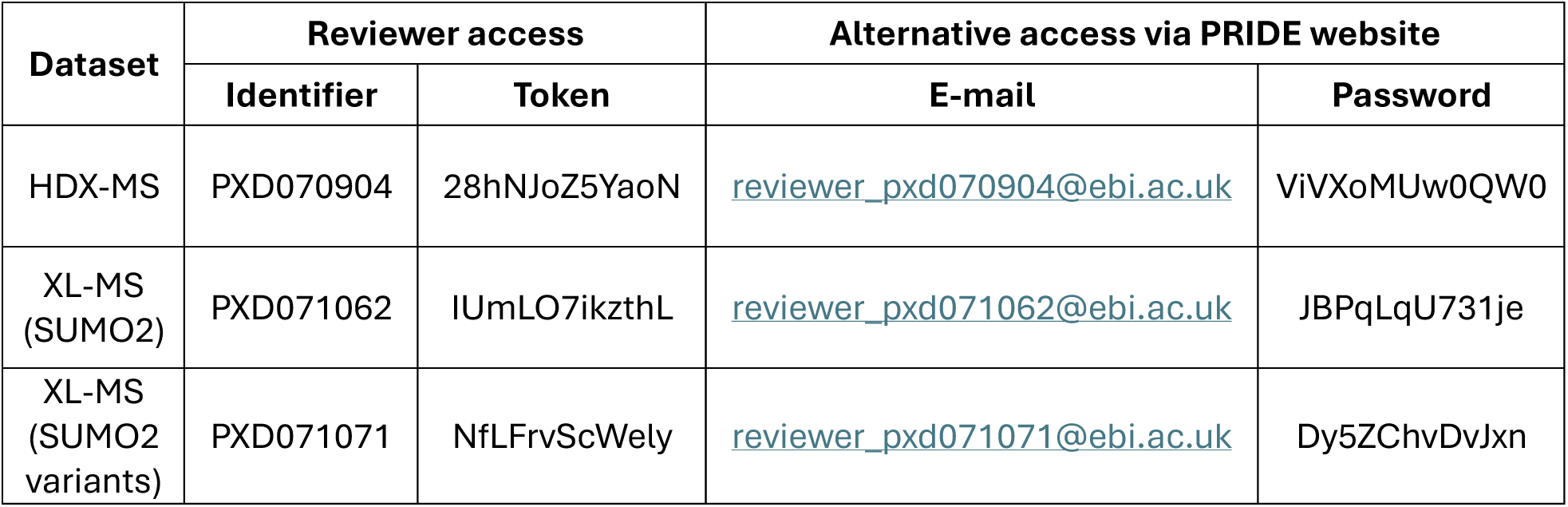

---

**Fig S1:**
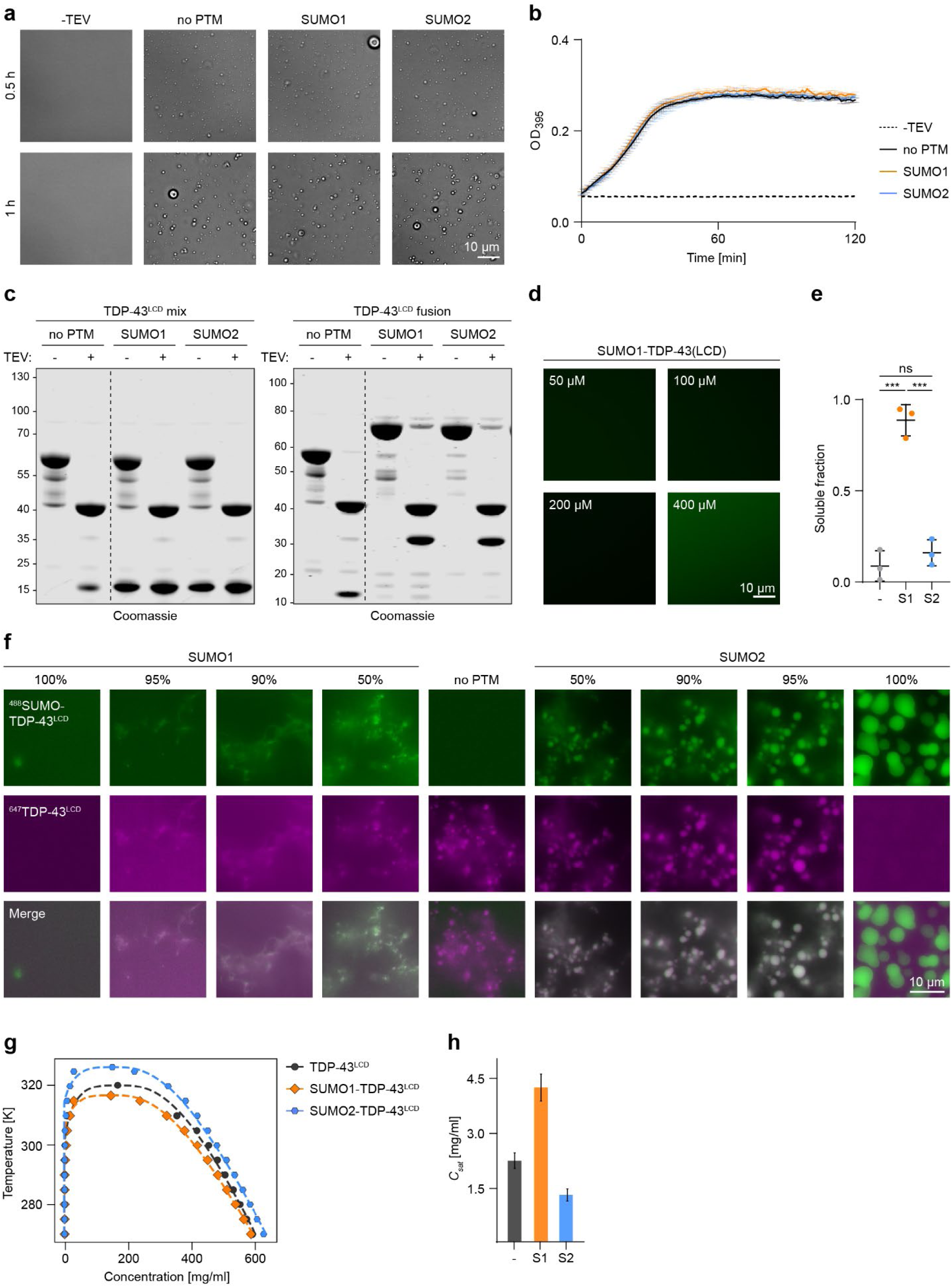
Covalent attachment is needed for SUMO’s effect on TDP-43^LCD^. **a-b,** Free SUMO does not affect phase separation of TDP-43^LCD^. **a,** Condensate formation of 50 µM TDP-43^LCD^ mixed with 50 µM SUMO as indicated, monitored by DIC microscopy at indicated times after addition of TEV protease. **b,** Turbidity measurement of 50 µM TDP-43^LCD^ mixed with 50 µM SUMO as indicated, after addition of TEV protease. Values represent means ±SD of 3 technical replicates (one of three independent experiments). **c,** Confirmation of TEV-induced MBP cleavage from TDP-43^LCD^ constructs by SDS-PAGE and Coomassie staining, performed at the end of the assays shown in panel b and Fig. 1c. **d,** Fusion of SUMO1 inhibits TDP-43^LCD^ phase separation at up to 400 µM protein concentration. Condensate formation of SUMO1-TDP-43^LCD^ at the indicated concentrations (5% AzDye488-labelled), monitored 1 h after addition of TEV protease. **e,** Quantification of the sedimentation assay shown in Fig. 1d by band intensities as a measure of solubility (Sn/(Sn+Pt) from three independent experiments (means ±SD, two-tailed paired Student’s t-test, * *p*<0.05; ** *p*<0.005; ns: not significant). **f,** Fusion of SUMO affects TDP-43^LCD^ phase separation at substoichiometric concentration. Fluorescence microscopy of defined mixtures of SUMO-TDP-43^LCD^ (5% AzDye488-labelled) and unmodified TDP-43^LCD^ (5% Atto647-labelled) at a total protein concentration of 50 µM, monitored 1 h after the addition of TEV protease. **g,** Phase diagram of the simulated systems, showing the coexistence between dilute and dense phases as a function of temperature and overall concentration. Symbols indicate coexistence points obtained from direct coexistence simulations, and lines are fit to the coexistence data using the functional form for the three-dimensional Ising universality class^92^. **h,** Mean saturation concentrations of the dilute phases at 300 K for all simulated systems, extracted from the coexistence data shown in panel g. Error bars indicate SDs across three independent simulation replicas.

**Fig S2:**
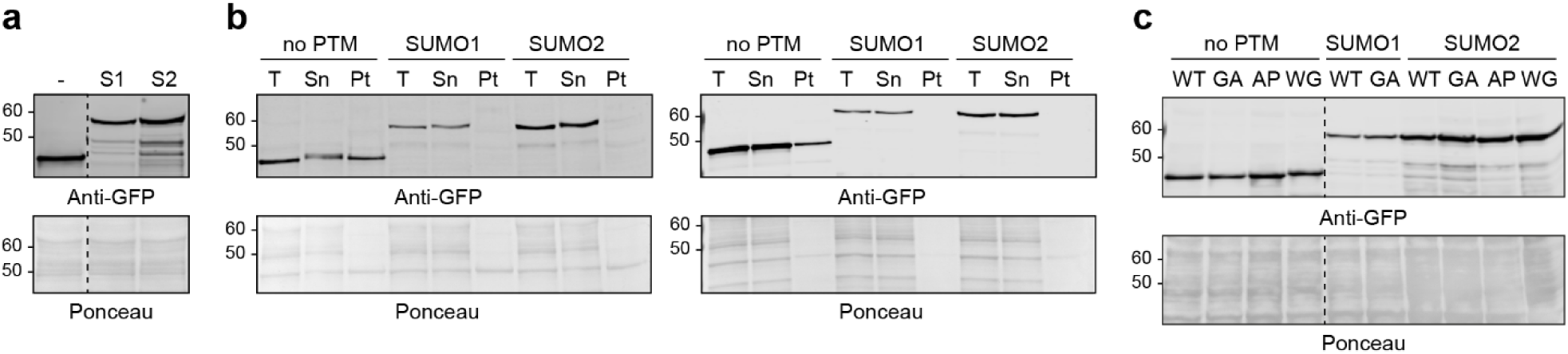
TDP-43^LCD^-eGFP expression and sedimentation profiles. **a,** Expression levels of TDP-43^LCD^-eGFP constructs, analysed by SDS-PAGE and western blotting using an anti-GFP antibody. Ponceau staining served as loading control. Dashed lines indicate removal of irrelevant portions from the blot. **b,** Two independent replicates of the experiment shown in Fig. 2f-g. **c,** Expression levels of TDP-43^LCD^-eGFP phase separation mutants, analysed by SDS-PAGE and western blotting. Ponceau staining served as loading control. Dashed lines indicate removal of irrelevant portions from the blot.

**Fig S3:**
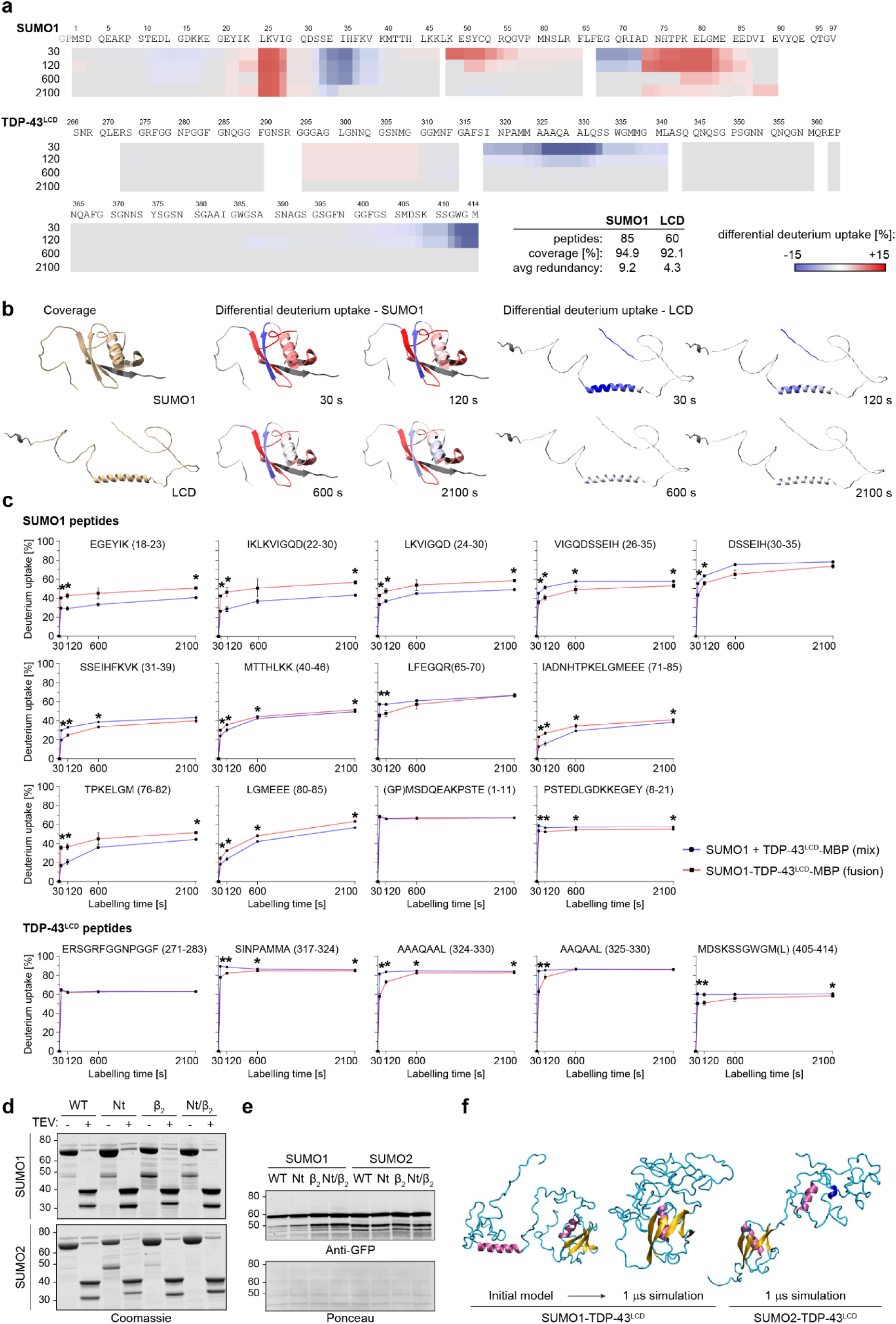
HDX-MS reveals intramolecular contacts between SUMO1 and TDP-43^LCD^. **a,** Heat map presentation of differential deuterium uptake after labelling for 30, 120, 600 and 2,100 s, plotted onto the sequences of SUMO1 and TDP-43^LCD^. **b,** Peptide coverage (in gold) and differential deuterium uptake after 30, 120, 600 and 2,100 s labelling, plotted onto AlphaFold 3 models of SUMO1 and TDP-43^LCD^. Residues not identified by MS are shown in dark grey. **c,** Relative deuterium uptake plots of selected peptides derived from SUMO1 or TDP-43^LCD^, comparing a non-covalent mix of SUMO1 and TDP-43^LCD^ to the SUMO1-TDP-43^LCD^ fusion protein. **d,** Confirmation of TEV-induced cleavage of MBP from SUMO-TDP-43^LCD^ hybrid proteins by SDS-PAGE and Coomassie staining at the end of the assay shown in Fig. 3d. **e,** Expression levels of SUMO-TDP-43^LCD^-eGFP hybrid proteins analysed by SDS-PAGE and western blotting using an anti-GFP antibody. Ponceau staining served as loading controls. **f,** One of the initial atomistic models of the SUMO1-TDP-43^LCD^ used as starting structure for MD simulations, along with representative snapshots of the SUMO1-TDP-43^LCD^ systems after 1 μs of simulation, in which the CR region of TDP-43LCD is completely disrupted, and the SUMO2-TDP-43^LCD^ system, which adopts a more extended conformation (magenta and blue: α-helices; yellow: β-sheet; cyan: disordered).

**Fig S4:**
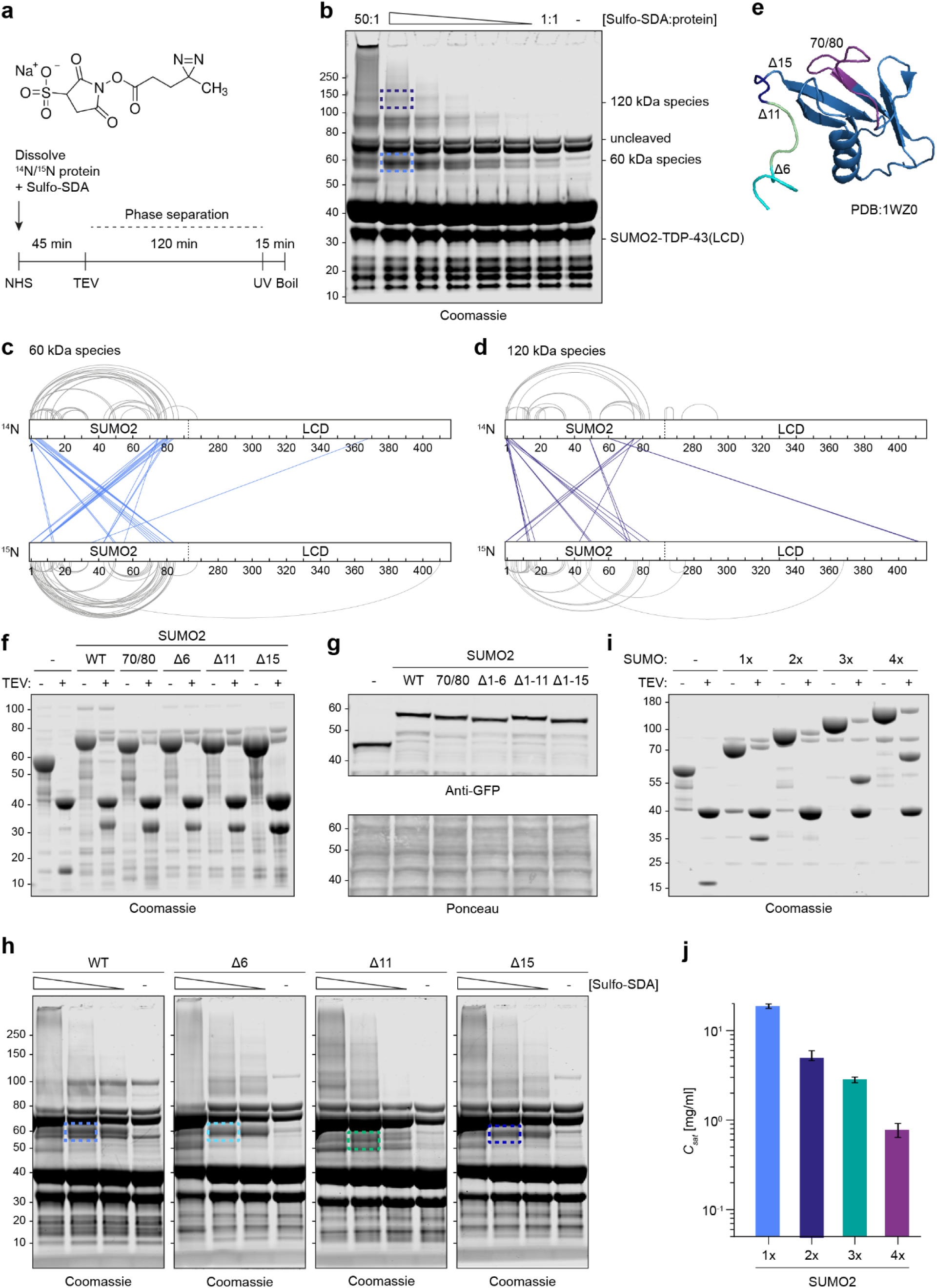
XL-MS and MD simulations reveal intermolecular self-association of SUMO2. **a,** Structure of the bifunctional crosslinker, sulfosuccinimidyl 4,4’-azipentanoate (Sulfo-SDA, top) and workflow of the crosslinking reaction under phase separation conditions. **b,** Titration of the Sulfo-SDA crosslinker in a reaction with a 1:1 mixture of ^14^N- and ^15^N-SUMO2-TDP-43^LCD^, analysed by SDS-PAGE and Coomassie staining. Gel fragments excised for LC-MS analysis are indicated by boxes. **c-d,** *Cis*- and *trans*-crosslinks plotted onto SUMO2-TDP-43^LCD^ for the 60 kDa (c) and 120 kDa species (d). Amino acids are numbered according to the original proteins. **e,** Structure of SUMO2 with *trans*-crosslinked regions selected for mutagenesis (N-terminus and the 70/80-loop) highlighted in different colours. **f,** Confirmation of TEV-induced cleavage of MBP from SUMO2-TDP-43^LCD^ interaction mutants by SDS-PAGE and Coomassie staining at the end of the assay shown in Fig. 4d. **g,** Expression levels of SUMO2-TDP-43^LCD^-eGFP interaction mutants analysed by SDS-PAGE and western blotting using an anti-GFP antibody. Ponceau staining served as loading control. **h,** Titration of the Sulfo-SDA crosslinker in reaction with 1:1 mixtures of ^14^N- and ^15^N-SUMO2-TDP-43^LCD^ truncation mutants, analysed by SDS-PAGE and Coomassie staining. Gel fragments excised for LC-MS analysis are indicated by boxes. **i,** Analysis of TEV-induced cleavage of MBP from poly-SUMO2-TDP-43^LCD^ proteins by SDS-PAGE and Coomassie staining. **i,** Quantification of saturation concentrations (*C_sat_*) corresponding to the MD simulations of poly-SUMO2-TDP-43^LCD^ proteins shown in Fig 4j.

**Fig S5:**
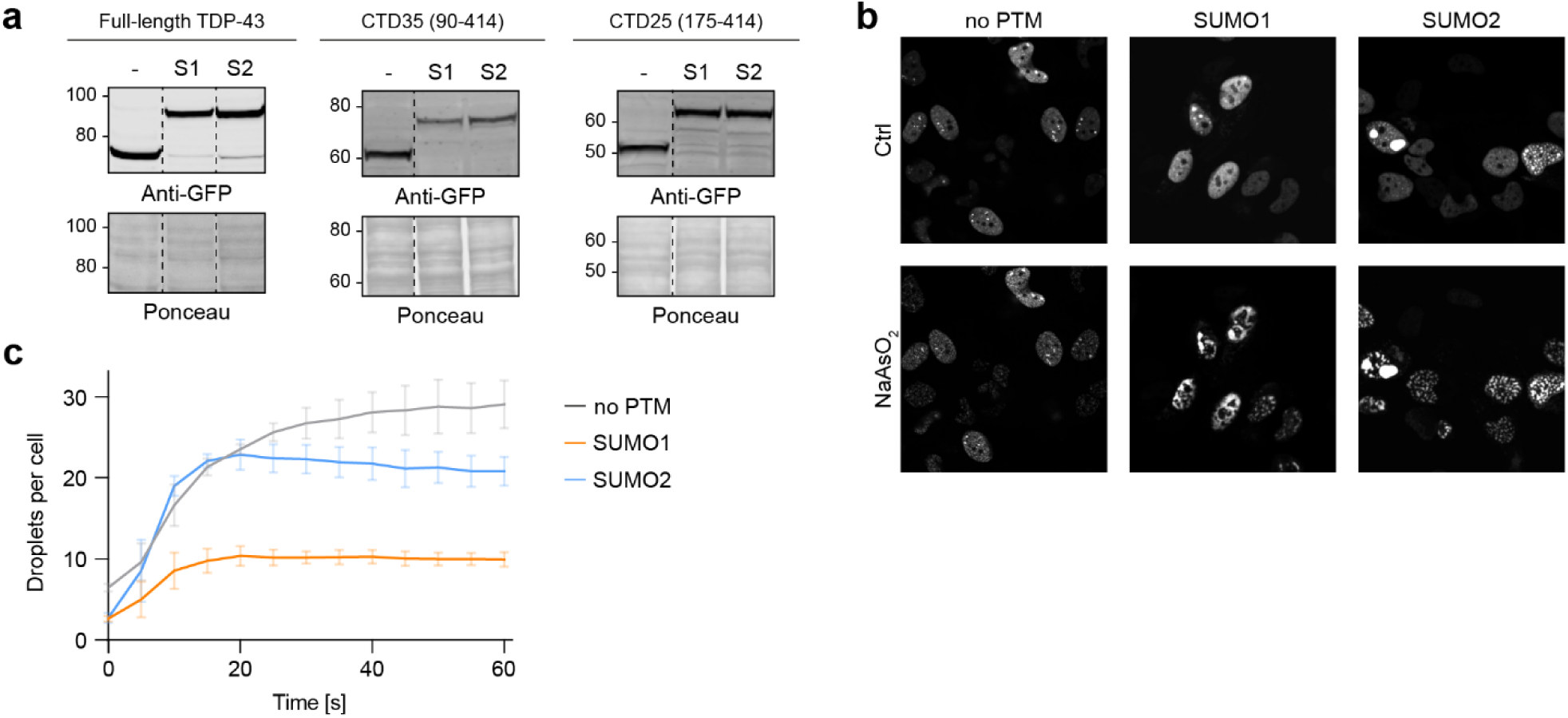
SUMO isoforms modulate phase separation of cellular TDP-43-eGFP. **a,** Expression levels of the constructs used in Fig. 5a-d, analysed by SDS-PAGE and western blotting using an anti-GFP antibody. Ponceau staining served as loading control. Dashed lines indicate removal of irrelevant portions from the blot. **b,** Modification with SUMO1 reduces stress-induced condensate formation of TDP-43. Representative images of the experiment shown in Fig. 5f. HeLa cells expressing full-length TDP-43-eGFP fused to SUMO isoforms were exposed to oxidative stress (100 µM NaAs_2_) for 60 min and subjected to fluorescence microscopy. **c,** Quantification of TDP-43-eGFP nuclear bodies (average absolute numbers ±SD) after exposure to oxidative stress as in panel b, calculated from three independent biological replicates.

**Fig S6:**
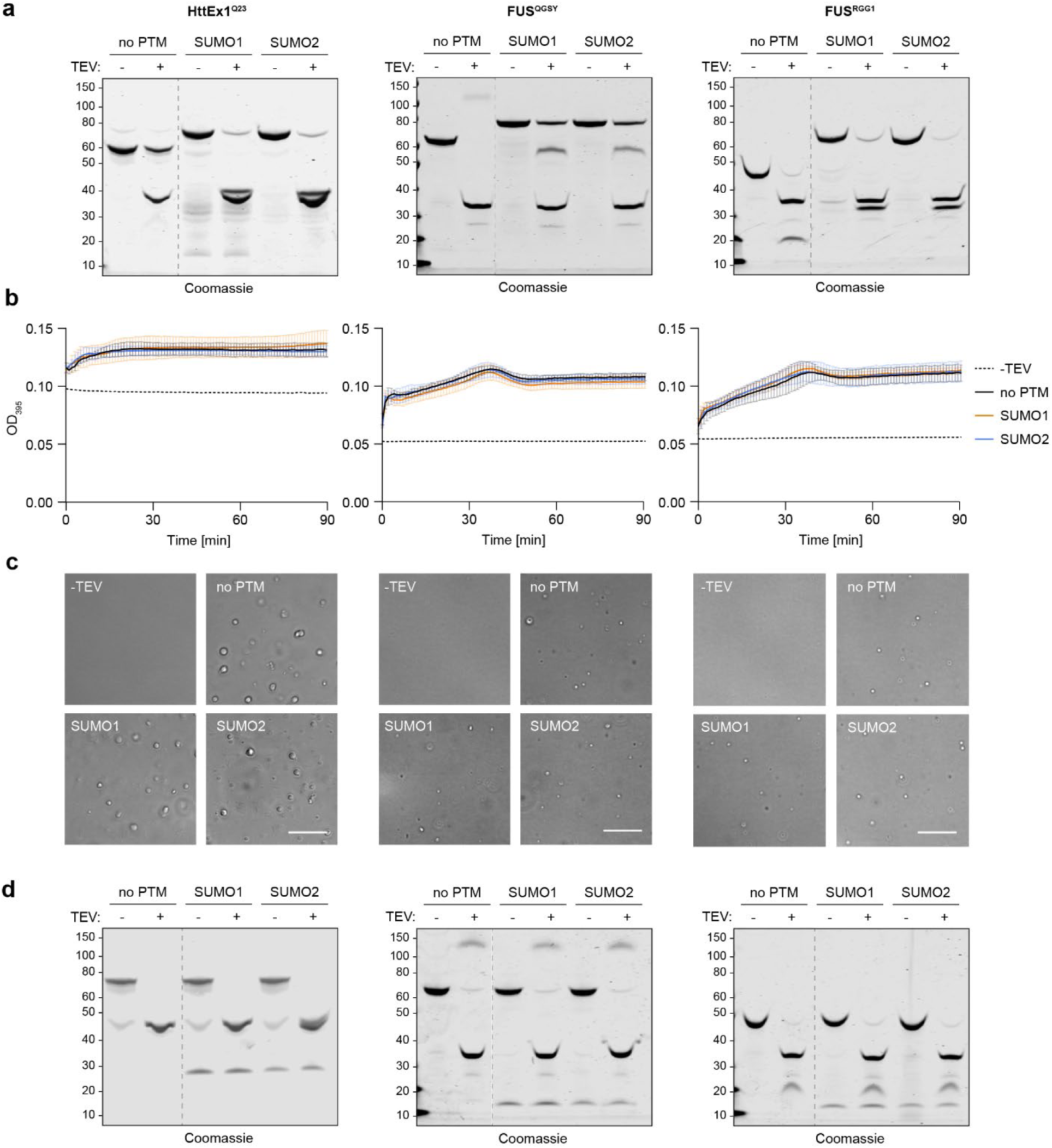
Covalent attachment of SUMO is required for its effect on other IDRs. **a,** Analysis of TEV-induced MBP-cleavage from HttEx1^Q23^ (left), FUS^QGSY^ (middle), and FUS^RGG1^ (right) fused to SUMO isoforms. **b-d,** Addition of free SUMO has no effect on the phase separation of Huntingtin- and FUS-derived IDRs. **a,** Turbidity measurements of the indicated domains mixed with equimolar amounts of SUMO isoforms, after addition of TEV protease (HttEx1^Q23^: 20 µM in 10% (w/v) PEG-8,000; FUS^QGSY^ and FUS^RGG1^: 10 µM in 5% (w/v) PEG-8,000). Values represent means ±SD of 3 technical replicates (one out of three measurements). TEV cleavage patterns are shown in panel d. **b,** DIC microscopy of HttEx1^Q23^ (left), FUS^QGSY^ (middle) and FUS^RGG1^ (right) in 1:1 mixtures with SUMO isoforms, analysed 1 h after cleavage of MBP by TEV protease under similar conditions as in panel a. **c,** Analysis of TEV-induced MBP-cleavage from HttEx1^Q23^ (left), FUS^QGSY^ (middle) and FUS^RGG1^ (right) mixed with free SUMO. **d,** Analysis of TEV-induced MBP-cleavage from HttEx1^Q23^ (left), FUS^QGSY^ (middle), and FUS^RGG1^ (right), mixed with SUMO isoforms.

## Supplementary Material

**Table S1.** Plasmids used in this study.

| ID | Name | Selection |
| --- | --- | --- |
| ***E. coli* expression vectors for recombinant protein production** |  |  |
| 5134 | pET15-His-FKBP-3C-2xSUMO2-TDP-43(LCD)-TEV-MBP | Amp |
| 5277 | pET15-His-FKBP-3C-TDP-43(LCD, G275C)-TEV-MBP | Amp |
| 5377 | pET15-His-FKBP-3C-4xSUMO2-TDP-43(LCD)-TEV-MBP | Amp |
| 5406 | pET15-His-FKBP-3C-SUMO2-TDP-43(LCD)-SUMO2(Δ1-10)-TEV-MBP | Amp |
| 5419 | pET15-His-FKBP-L20-3C-FUS(QGSY)-TEV-MBP | Amp |
| 5473 | pET15-His-FKBP-L20-3C-SUMO1-FUS(QGSY)-TEV-MBP | Amp |
| 5474 | pET15-His-FKBP-L20-3C-SUMO2-FUS(QGSY)-TEV-MBP | Amp |
| 5513 | pET15-His-FKBP-L20-3C-FUS(RGG)-TEV-MBP | Amp |
| 5515 | pET15-His-FKBP-L20-3C-SUMO1-FUS(RGG)-TEV-MBP | Amp |
| 5516 | pET15-His-FKBP-L20-3C-SUMO2-FUS(RGG)-TEV-MBP | Amp |
| 5517 | pET15-His-FKBP-L20-3C-SUMO1-TDP-43(LCD)-TEV-MBP | Amp |
| 5518 | pET15-His-FKBP-L20-3C-SUMO2-TDP-43(LCD)-TEV-MBP | Amp |
| 5542 | pCoofy1-His-3C-SUMO1 | Kan |
| 5543 | pCoofy1-His-3C-SUMO2 | Kan |
| 5582 | pET15-His-FKBP-L20-3C-Htt-Ex1-Q23-TEV-MBP | Amp |
| 5636 | pET15-His-FKBP-L20-3C-SUMO1-Htt-Ex1-Q23-TEV-MBP | Amp |
| 5637 | pET15-His-FKBP-L20-3C-SUMO2-Htt-Ex1-Q23-TEV-MBP | Amp |
| 5724 | pET15-His-FKBP-L20-3C-SUMO1-TDP-43(LCD, G275C)-TEV-MBP | Amp |
| 5725 | pET15-His-FKBP-L20-3C-SUMO2-TDP-43(LCD, G275C)-TEV-MBP | Amp |
| 5959 | pET15-His-FKBP-L20-3C-TDP-43(LCD)-TEV-MBP | Amp |
| 5982 | pET15-His-FKBP-L20-3C-SUMO1(SUMO2(1-15))-TDP-43(LCD)-TEV-MBP | Amp |
| 5985 | pET15-His-FKBP-L20-3C-SUMO2(SUMO1(1-19))-TDP-43(LCD)-TEV-MBP | Amp |
| 6072 | pET15-His-FKBP-L20-3C-SUMO1(SUMO2(29-35))-TDP-43(LCD)-TEV-MBP | Amp |
| 6073 | pET15-His-FKBP-L20-3C-SUMO1(SUMO2(1-15, 29-35))-TDP-43(LCD)-TEV-MBP | Amp |
| 6074 | pET15-His-FKBP-L20-3C-SUMO2(SUMO1(33-39))-TDP-43(LCD)-TEV-MBP | Amp |
| 6075 | pET15-His-FKBP-L20-3C-SUMO2(SUMO1(1-19,33-39))-TDP-43(LCD)-TEV-MBP | Amp |
| 6399 | pET15-His-FKBP-L20-3C-SUMO2(Δ1-6)-TDP-43(LCD)-TEV-MBP | Amp |
| 6400 | pET15-His-FKBP-L20-3C-SUMO2(Δ1-11)-TDP-43(LCD)-TEV-MBP | Amp |
| 6505 | pET15-His-FKBP-L20-3C-SUMO2(70/80-neutral)-TDP-43(LCD)-TEV-MBP | Amp |

| **Mammalian expression vectors** |  |  |
| --- | --- | --- |
| 5685 | pCMV-His8-TDP-43(LCD)-EGFP | Kan |
| 5688 | pCMV-His8-SUMO1-TDP-43(LCD)-EGFP | Kan |
| 5689 | pCMV-His8-SUMO2-TDP-43(LCD)-EGFP | Kan |
| 5936 | pCMV-His8-TDP-43-EGFP | Kan |
| 5938 | pCMV-His8-SUMO1-TDP-43-EGFP | Kan |
| 5639 | pCMV-His8-SUMO2-TDP-43-EGFP | Kan |
| 5988 | pCMV-His8-TDP-43(LCD,G335A,G338A)-EGFP | Kan |
| 5989 | pCMV-His8-SUMO1-TDP-43(LCD,G335A,G338A)-EGFP | Kan |
| 5990 | pCMV-His8-TDP-43(LCD,A326P)-EGFP | Kan |
| 5991 | pCMV-His8-SUMO2-TDP-43(LCD,A326P)-EGFP | Kan |
| 5992 | pCMV-His8-TDP-43(LCD,Δ2W)-EGFP | Kan |
| 5993 | pCMV-His8-SUMO2-TDP-43(LCD,Δ2W)-EGFP | Kan |
| 6022 | pCMV-His8-SUMO2-TDP-43(LCD,G335A,G338A)-EGFP | Kan |
| 6056 | pCMV-His8-SUMO1(SUMO2(29-35))-TDP-43(LCD)-EGFP | Kan |
| 6057 | pCMV-His8-SUMO1(SUMO2(1-15,29-35))-TDP-43(LCD)-EGFP | Kan |
| 6058 | pCMV-His8-SUMO2(SUMO1(33-39))-TDP-43(LCD)-EGFP | Kan |
| 6059 | pCMV-His8-SUMO2(SUMO1(1-19,33-39))-TDP-43(LCD)-EGFP | Kan |
| 6493 | pCMV-His8-TDP-43(CTF25)-EGFP | Kan |
| 6494 | pCMV-His8-TDP-43(CTF35)-EGFP | Kan |
| 6514 | pCMV-His8-SUMO2(70/80-neutral)-TDP-43(LCD)-EGFP | Kan |
| 6545 | pCMV-His8-SUMO2(Δ1-6)-TDP-43(LCD)-EGFP | Kan |
| 6546 | pCMV-His8-SUMO2(Δ1-11)-TDP-43(LCD)-EGFP | Kan |
| 6547 | pCMV-His8-SUMO2(Δ1-15)-TDP-43(LCD)-EGFP | Kan |
| 6502 | pCMV-His8-SUMO1-TDP-43(CTF25)-EGFP | Kan |
| 6503 | pCMV-His8-SUMO2-TDP-43(CTF25)-EGFP | Kan |
| 6505 | pCMV-His8-SUMO1-TDP-43(CTF35)-EGFP | Kan |
| 6506 | pCMV-His8-SUMO1-TDP-43(CTF35)-EGFP | Kan |

